# p38α MAPK integrates opposing oncogenic APC/β-catenin/TCF and GUCY2C tumor-suppressor axes in colorectal cancer

**DOI:** 10.64898/2026.10.05.756833

**Authors:** Ariana A. Entezari, Adi Caspi, Jasmine R. Alvarez, Andrew E. Evans, Jeffrey A. Rappaport, Thomas J.M. Kuret, Allison S. Doermann, Ramkrishna Mitra, Adam E. Snook, Scott A. Waldman

## Abstract

**Introduction:** Sporadic colorectal cancers (CRCs) form following initiating APC mutations, inducing constitutive β-catenin/TCF signaling. Similarly, the guanylyl cyclase C (GUCY2C) tumor suppressor axis is universally silenced in CRC. While the GUCY2C receptor is retained, loss of its paracrine hormones guanylin (GUCA2A) and uroguanylin (GUCA2B) is among the earliest events in transformation. Mutant APC/β-catenin/TCF signaling driving CRC transcriptionally silences GUCY2C hormone expression while suppressing that oncogenic pathway reconstitutes hormone production opposing tumorigenesis. p38α opposes tumor initiation, and its loss promotes transformation in chemical and genetic mouse models of CRC. In healthy intestine, p38α supports a gradient of epithelial differentiation along the crypt-surface axis. Here, we reveal that p38α links regulation of oncogenic APC/β-catenin/TCF signaling and GUCY2C hormone expression.

**Methods:** Regulation of p38α by APC/β-catenin/TCF signaling was defined in four genetically engineered human CRC cell lines with conditional oncogenic APC/β-catenin/TCF signaling and healthy human colon organoids. Expression of p38α was eliminated from conditional human CRC cell lines by pharmacological inhibition, RNA knockdown, and CRISPR/Cas9 gene editing. Transcriptomic analysis of these cells by RNA sequencing defined the set of genes regulated by APC/β-catenin/TCF signaling controlled through p38α.

**Results:** In CRC cell lines, APC/β-catenin/TCF signaling suppressed p38α phosphorylation and activity, whereas inhibition of APC/β-catenin/TCF restored p38α signaling. In the absence of APC/β-catenin/TCF signaling, inhibiting p38α repressed GUCY2C hormone expression. Inhibiting APC/β-catenin/TCF signaling enriched phosphorylated p38α at the GUCA2A promoter. In that context, analysis of the CPTAC proteomic database revealed that phosphorylation of p38α and its upstream kinases MKK3 and MKK4 was reduced in human CRC. Finally, transcriptomic analysis revealed that p38α is a critical signaling node controlling the expression of a substantial subset of APC/β-catenin/TCF-regulated genes in human CRC cells and patient tumors.

**Conclusions:** APC/β-catenin/TCF signaling inhibits p38α phosphorylation and transcriptional activity, suppressing GUCA2A and GUCA2B expression and functionally silencing the retained GUCY2C receptor. These findings identify p38α as a signaling intermediary integrating oncogenic APC/β-catenin/TCF activity to extinction of the GUCY2C tumor-suppressor axis and reveal a broader p38α-dependent component of the APC/β-catenin/TCF-regulated transcriptome.

Graphical Abstract

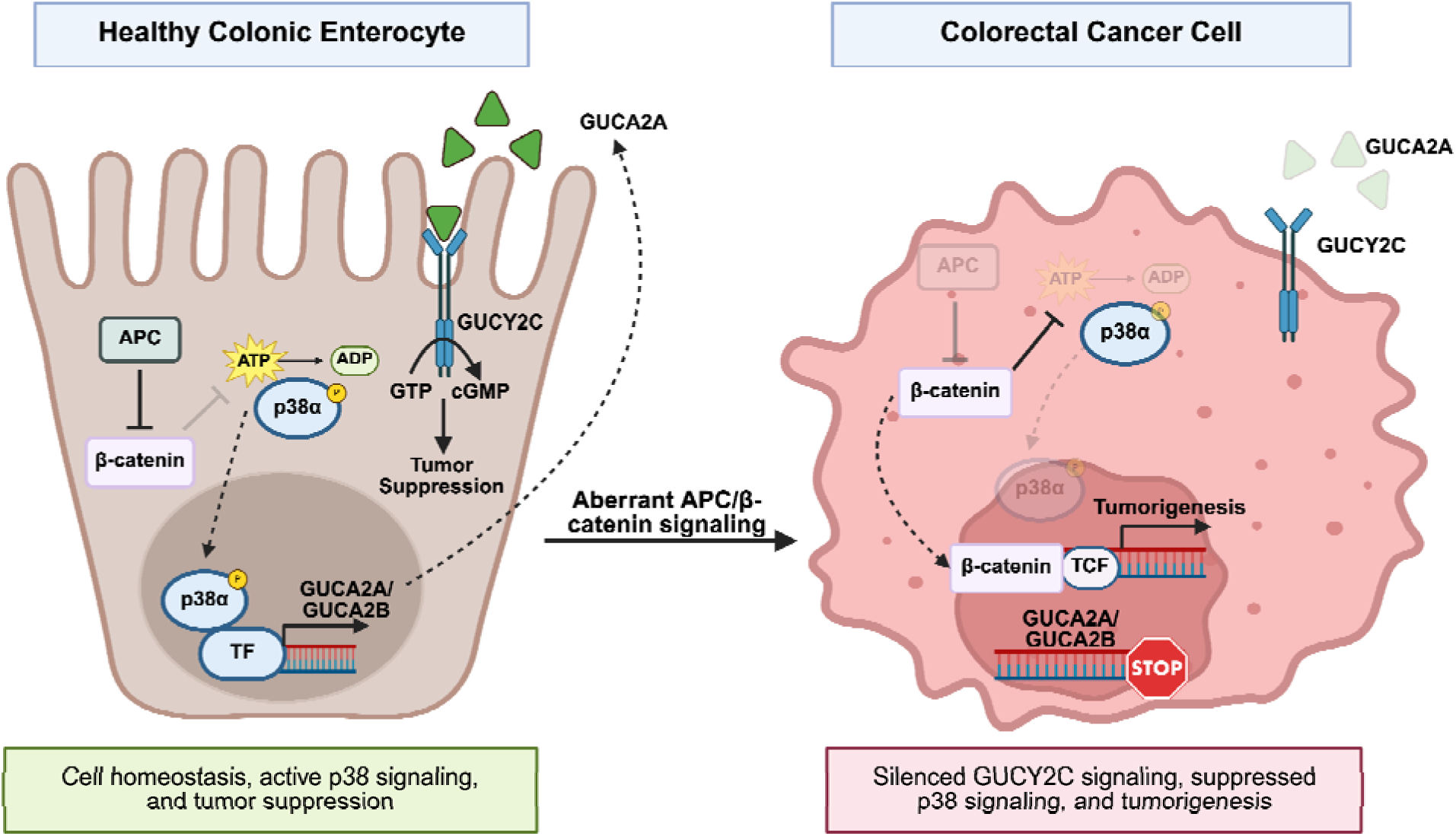

## Introduction

The Wnt downstream signaling pathway mediated by APC/β-catenin/TCF is dysregulated in most colorectal cancers (CRCs), acting as a driver for tumor development and progression.^1^ Gain-of-function mutations in this oncogenic pathway drive nuclear translocation of β-catenin which binds TCF transcription factors activating the expression of genes such as *MYC*, *LGR5*, and cyclins that promote transformation.^1^ Additionally, a substantial fraction of targets are transcriptionally repressed by this oncogenic pathway.^2,3^ While activated gene programs are well-characterized, the regulatory architecture governing repressed genes remains under-explored.^1,3^

One-third of genes repressed by gain-of-function mutations in Wnt signaling in colorectal and pancreatic tumors appear to be mediated by the ERK MAPK pathway^3^ which coordinates fundamental processes including proliferation, survival, and apoptosis.^4^ Similarly, p38 is a MAPK signaling node comprising α, β, γ, and δ isoforms.^5^ In the intestine, phosphorylation of p38α and p38δ regulates cell proliferation and differentiation, intestinal epithelial barrier function, and immune responses.^5–7^ Dysregulation of intestinal p38 signaling is associated with CRC initiation,^5,6,8^ reflecting its role in regulating cell cycle arrest and apoptosis.^9,10^ A linkage between APC/β-catenin/TCF and p38 MAPK pathways in CRC has not been defined previously.

The epithelial tumor suppressor guanylyl cyclase C (GUCY2C) is a transmembrane receptor activated by endogenous hormones guanylin (GUCA2A) and uroguanylin (GUCA2B) in the large and small intestine, respectively.^11–14^ Activation of GUCY2C drives cGMP accumulation and secretion of fluid and electrolytes across the mucosa.^14^ Also, GUCY2C controls homeostatic processes dysregulated in colorectal tumorigenesis, including proliferation, metabolism, DNA repair, and differentiation.^15–20^ Eliminating GUCY2C expression in mice produces crypt hyperplasia, cell cycle acceleration, and DNA damage. Conversely, GUCY2C stimulation delays cell cycle progression, reduces DNA damage, and induces senescence.^15,16,18,19^ Importantly, GUCY2C signaling opposes APC/β-catenin/TCF-dependent intestinal tumorigenesis in mice.^15,16,20^ Universal silencing of GUCY2C signaling through loss of GUCA2A and GUCA2B occurs early in CRC and is conserved across tumor subtypes and species.^21,22^ Also, GUCA2A and GUCA2B expression is lost following conditional APC inactivation in mice.^23^ Further, silencing mutant APC/β-catenin/TCF signaling restores GUCA2A and GUCA2B mRNA and protein levels.^2,24^ Indeed, expression of GUCA2A and GUCA2B, and consequently GUCY2C signaling, is tightly regulated at the transcriptional level by the APC/β-catenin/TCF pathway.^2,24^

Our model suggests that there is a reciprocal regulatory circuit between GUCY2C and APC/β-catenin/TCF axes that is corrupted in CRC. In that model, the earliest mutations in the APC/β-catenin/TCF pathway silence GUCY2C signaling by suppressing hormone transcription. Conversely, reconstituting GUCY2C activity, including oral hormone replacement in preclinical models, opposes APC/β-catenin/TCF-driven intestinal tumorigenesis. However, the precise molecular mechanisms linking these reciprocally regulating oncogenic and tumor suppressing pathways remain unknown. Recently, we identified an APC/β-catenin/TCF-sensitive locus control region (LCR) in the region 5’ upstream of the GUCA2A coding sequence that regulates transcription of GUCA2A and GUCA2B.^2^ Here, we identify p38α as a molecular link integrating APC/β-catenin/TCF signaling with the GUCY2C tumor-suppressor axis. Indeed, p38α is necessary for GUCY2C hormone expression, and APC/β-catenin/TCF signaling reduces p38α phosphorylation and activity. Activated p38α is enriched at the GUCA2A regulatory locus, where it is required for LCR activity. Beyond GUCY2C hormones, we demonstrate that p38α is required for changes in expression of a substantial subset of genes regulated by oncogenic APC/β-catenin/TCF signaling in CRC.

## Results

### ERK signaling is not necessary for GUCY2C hormone expression

Since ERK signaling regulates up to one-third of the APC/β-catenin/TCF-repressed transcriptome,^3^ we sought to determine whether it is required for GUCY2C hormone expression. The FDA-approved MEK1/2 inhibitor trametinib was used in DLD1 cells expressing an inducible dominant negative TCF7L2 (DNTCF) transgene.^25^ Induction of DNTCF by DOX^2,24^ induced GUCA2A expression, consistent with its repression by mutant APC/β-catenin/TCF (Supplemental Fig. 1A). Trametinib concentrations sufficient to inhibit ERK1/2 phosphorylation (Supplemental Fig. 1A) had no effect on GUCA2A expression (Supplemental Fig. 1B). However, at higher concentrations, trametinib suppressed GUCA2A expression (Supplemental Fig. 1B). Previous phosphoproteomic analyses revealed that these trametinib concentrations produced off-target inhibition of p38α signaling.^26^

### Regulation of GUCY2C hormone expression by APC/β-catenin/TCF signaling is mediated by p38α

Signaling by p38 was modulated in human CRC cell lines engineered with conditional regulation of oncogenic APC/β-catenin/TCF signaling. Inducing a wild-type APC transgene using a zinc-inducible promoter in HT29 cells harboring mutant APC degrades β-catenin and disrupts β-catenin/TCF signaling (Fig. 1A).^27^ Inducing a short hairpin RNA (shRNA) targeting β-catenin using a DOX-inducible promoter in LS174T cells harboring a gain-of function mutation in β-catenin reduces its accumulation (Fig. 1A).^25,28^ Finally, inducing a dominant negative TCF7L2 (DNTCF) transgene using a DOX-inducible promoter in LS174T cells silences APC/β-catenin/TCF signaling (Fig. 1A).^2,24^ Pharmacologic inhibition of p38α/β with SB202190, quantified by loss of phosphorylation of the downstream target HSP27 (Fig. 1B, C),^29,30^ blocked GUCA2A and GUCA2B mRNA expression induced by silencing APC/β-catenin/TCF signaling in HT29 (APC) cells (Fig. 1D), LS174T (siBC) cells (Fig. 1E), and LS174T (DNTCF) cells (Supplemental Fig. 2A). SB203580, a specific p38α inhibitor, also blocked GUCA2A and GUCA2B mRNA expression in LS174T cells induced by DNTCF (Supplemental Fig. 2B) or siBC (Fig. 1F). Beyond CRC, we assessed the impact of p38 inhibition in healthy human colonoids cultured in the presence or absence of Wnt3a, to toggle wild type APC/β-catenin/TCF signaling on and off, respectively.^31^ Depriving colonoids of Wnt increased p-HSP27 levels, reflecting activation of p38 signaling, which was suppressed by SB202190 (Fig. 1G). Consistent with the restriction of GUCA2A expression to differentiated epithelium,^32–34^ Wnt-deprived colonoids expressed GUCA2A mRNA which was suppressed by SB202190 (Fig. 1H).

**Figure 1.**
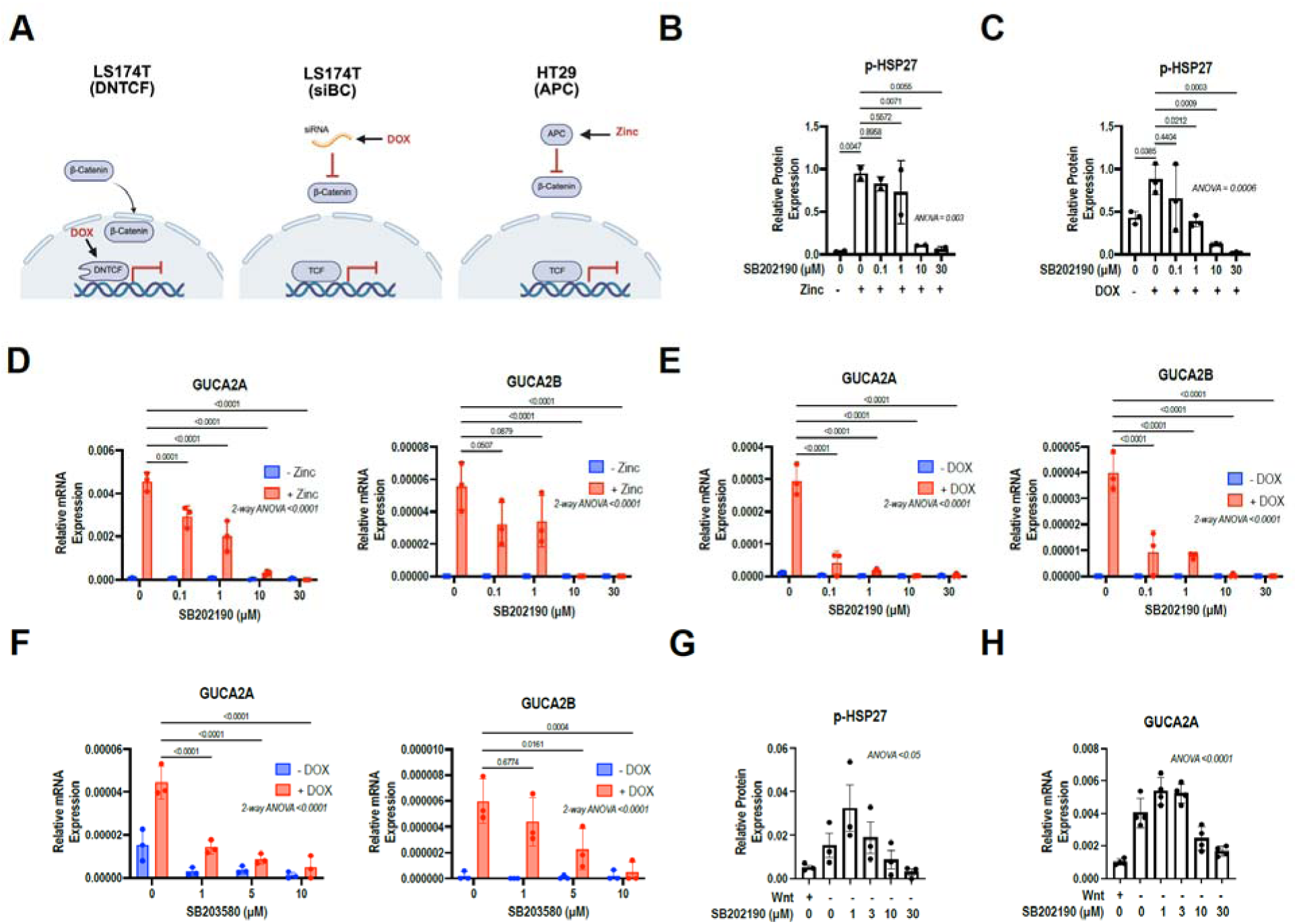
p38α/β inhibitors suppress mRNA expression of GUCA2A and GUCA2B in human CRC cells. (A) CRC cells express 1 of 3 inducible inhibitors of Wnt/β-catenin/TCF signaling: DOX-inducible dominant negative TCF7L2 (DNTCF; DLD1 and LS174T cells), DOX-inducible β-catenin siRNA (LS174T cells), or zinc-inducible wild-type APC (HT29 cells). (B,C) Phosphorylation of p38 target HSP27 was decreased in response to increasing concentrations of SB202190 ranging from 0-30 µM in (B) HT29 (APC) and (C) LS174T (siBC) cell lines. (D,E) GUCA2A and GUCA2B mRNA expression was decreased in response to increasing concentrations of SB202190 ranging from 0-30 µM in (D) HT29 (APC) and (E) LS174T (siBC) cell lines. (F) GUCA2A and GUCA2B mRNA expression was decreased in response to increasing concentrations of p38α inhibitor SB203580 in LS174T (siBC) cell lines. (G) Phosphorylated HSP27 protein and (H) GUCA2A mRNA increased in healthy human colonoids in response to loss of Wnt-conditioned medium. Under Wnt-deprived conditions, pHSP27 (G) and GUCA2A mRNA (H) were decreased after SB202190 treatment for 48h in human differentiated colon organoids. Statistics were calculated using a one-way ANOVA with Tukey’s multiple comparisons test (B, C, G, H) or a two-way ANOVA with Tukey’s multiple comparisons test (D-F). Error bars represent the standard deviation (SD).

Genetic knockdown of p38α in LS174T (DNTCF) cells using siRNA reduced GUCA2A expression following suppression of APC/β-catenin/TCF signaling (Supplemental Fig. 2C, D). Further, CRISPR/Cas9 knockout of p38α in LS174T (siBC), LS174T (DNTCF) and HT29 (APC) cells (Fig. 2A, B, Supplemental Fig. 3A-D), but not knockout of p38β, γ and δ expression in LS174T (siBC) cells (Fig. 2C, D), inhibited the induction of GUCA2A and GUCA2B mRNA expression by suppressing APC/β-catenin/TCF signaling. Moreover, re-expression of exogenous p38α in LS174T (DNTCF) p38α KO cells rescued GUCA2A and GUCA2B mRNA expression induced by silencing APC/β-catenin/TCF signaling (Fig. 2E, F).

**Figure 2.**
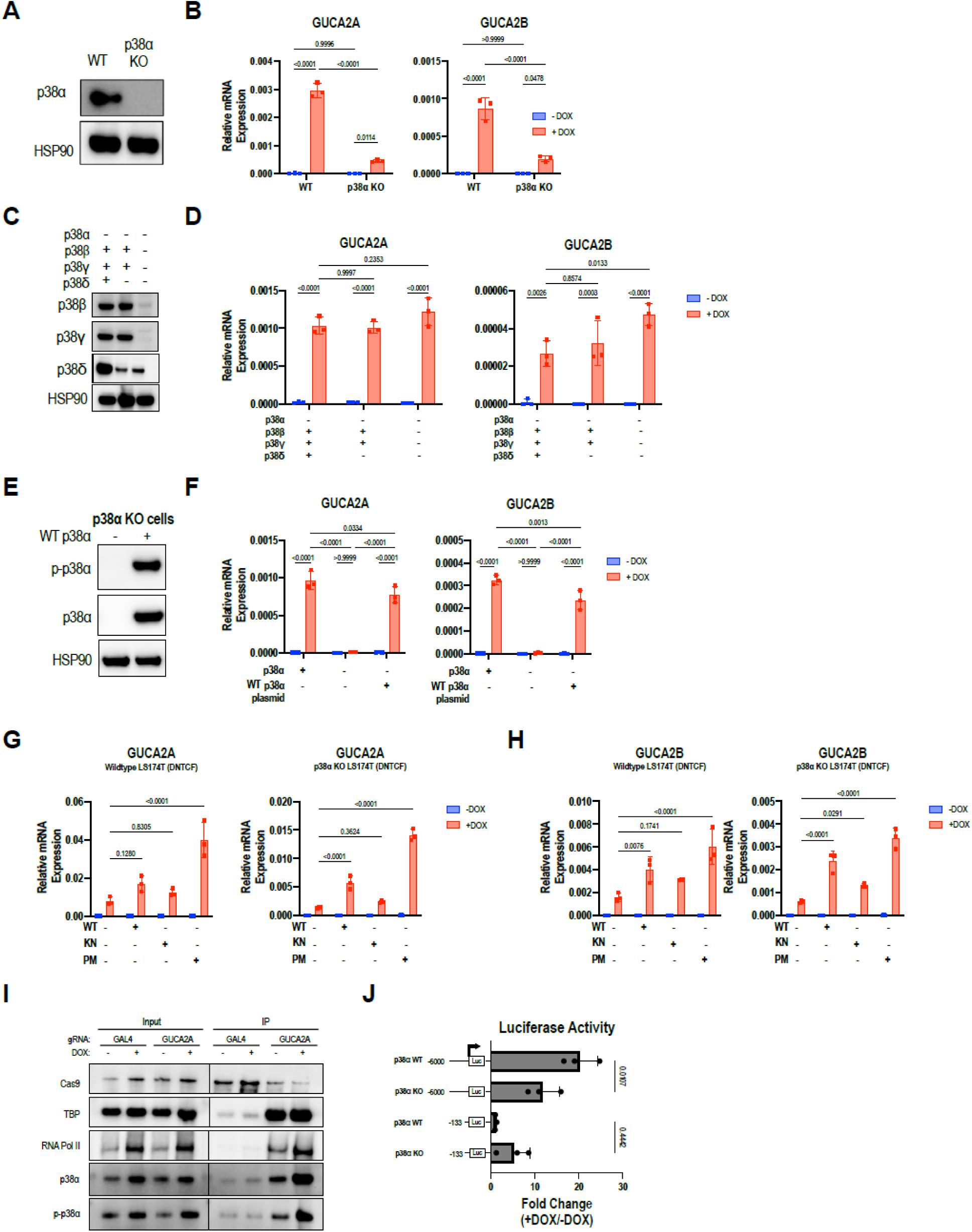
p38α CRISPR/Cas9 knockout reduces GUCA2A and GUCA2B mRNA expression in human CRC cells. (A) p38α is knocked out in LS174T (DNTCF) cells. (B) GUCA2A and GUCA2B mRNA expression is reduced in polyclonal p38α KO populations in LS174T (dnTCF) cells. (C) P38b, p38g, and p38d were knocked down in p38α KO LS174T (siBC) cells. (D) GUCA2A and GUCA2B mRNA expression is unchanged in p38α KO, p38d KD cells and p38α KO, p38b/p38g/p38d KD cells compared to p38α KO control cells. (E) p38α plasmid was transfected into single clonal LS174T (DNTCF) p38α KO cells. (F) WT p38α plasmid rescued GUCA2A and GUCA2B mRNA expression in single clonal LS174T (DNTCF) p38α KO cells. (G) GUCA2A and (H) GUCA2B mRNA expression is rescued by wildtype (WT) and phosphomimetic (PM) p38α, while an inactive kinase null (KN) construct is unable to upregulate expression. (I) CAPTURE followed by immunoblot in DLD1 (DNTCF) cells demonstrates enrichment of p38α and p-p38α at the GUCA2A promoter with a *GUCA2A*-targeting gRNA relative to a nontargeting gRNA (GAL4), with increased abundance +DOX (with DNTCF expression). The chromatin associated proteins RNA polymerase II and TATA-binding protein (TBP) serve as controls for chromatin pulldown. (J) Luciferase constructs driven by the indicated GUCA2A DNA regions were expressed in L8 (DNTCF) cells. Luciferase activity was quantified with (+) or without (–) 1 μg/mL DOX for 24 h, and data are presented relative to noninduced cells. Luciferase constructs were driven by the GUCA2A promoter (+15 to −133) or enhancer (+15 to –6000) relative to the GUCA2A transcription start site. Statistics were calculated using a two-way ANOVA with Tukey’s multiple comparisons test (B, D, F-I). Error bars represent the standard deviation (SD).

To further characterize the contribution of p38α to GUCA2A and GUCA2B expression, we introduced overexpression constructs consisting of a constitutively inactive (kinase-null, KN) construct with mutated phosphorylation sites, and a constitutively active (phosphomimetic, PM) construct with previously characterized mutations at amino acids 176 and 327 (Supplemental Fig. 3E).^35,36^ In both wildtype and p38α KO cells, the inactive KN construct did not change GUCA2A and GUCA2B expression over baseline (Fig. 2G, H). In contrast, the constitutively active construct increased GUCA2A and GUCA2B expression in those cell types (Fig. 2G, H). Notably, these constructs did not rescue expression in the absence of DOX, suggesting that p38α is insufficient for GUCY2C hormone transcription in the presence of APC/β-catenin/TCF signaling, and that repression must be relieved through an additional mechanism in the context of active APC/β-catenin/TCF signaling. These findings indicate that p38α positively regulates GUCA2A/B expression through its kinase activity.

### p38α is enriched at the GUCA2A promoter in the absence of APC/β-catenin/TCF signaling

We employed <u>C</u>RISPR <u>a</u>ffinity <u>p</u>urification *in si<u>tu</u>* of regulatory <u>e</u>lements (CAPTURE) to identify proteins interacting with the *GUCA2A* gene locus.^37^ CAPTURE utilizes a catalytically dead (d)Cas9 protein containing a biotin acceptor site coupled to a targeting gRNA. In the presence of birA, a prokaryotic biotin ligase, dCas9 is biotinylated *in situ* and protein complexes surrounding dCas9 are chemically crosslinked. Finally, streptavidin purification isolates proteins associated with the captured locus (Supplemental Fig. 3F). A gRNA targeting the *GUCA2A* proximal promoter was validated through streptavidin ChIP to effectively capture the targeted locus (Supplemental Fig. 3G).^2^ CAPTURE analysis revealed enrichment of p38α and phosphorylated (activated) p38α (p-p38) at the *GUCA2A* locus following DNTCF inhibition of APC/β-catenin/TCF signaling (Fig. 2I).

### Expression of GUCY2C hormones by p38α is regulated by the LCR

GUCA2A and GUCA2B mRNA expression induced by silencing APC/β-catenin/TCF signaling is mediated by a LCR controlling transcription of both hormones, contained within the 6kb 5’ region of the GUCA2A transcriptional start site.^2^ In that context, DOX induction of DNTCF in LS174T cells inhibited APC/β-catenin/TCF signaling and restored GUCA2A and GUCA2B mRNA expression, associated with induction of luciferase expression produced by a reporter containing the 5’ GUCA2A LCR, and this was reduced ∼50% by knocking out p38α (Fig. 2J). This indicates that p38α regulates GUCA2A and GUCA2B through a mechanism requiring the Wnt-sensitive LCR.

### APC/β-catenin/TCF suppresses p38α signaling in CRC cells

Since p38α is necessary for de-repression of GUCY2C hormone expression after silencing APC/β-catenin/TCF signaling, this kinase may be inhibited by that oncogenic pathway which contributes to GUCA2A and GUCA2B loss. Previous work identified a set of p38α/β-regulated genes induced by TNFα, anisomycin, and NaCl in mouse embryonic fibroblasts.^38^ Gene set enrichment analysis (GSEA) in the four conditional cell lines revealed that silencing APC/β-catenin/TCF signaling induced this p38α/β signature, suggesting opposing roles for these pathways (Fig. 3A). Additionally, while total p38α protein levels remained unchanged (Fig. 3B), phosphorylation of p38α (T180/Y182; Fig. 3C) and its downstream target HSP27 (Fig. 3D) was inhibited by APC/β-catenin/TCF signaling.

**Figure 3.**
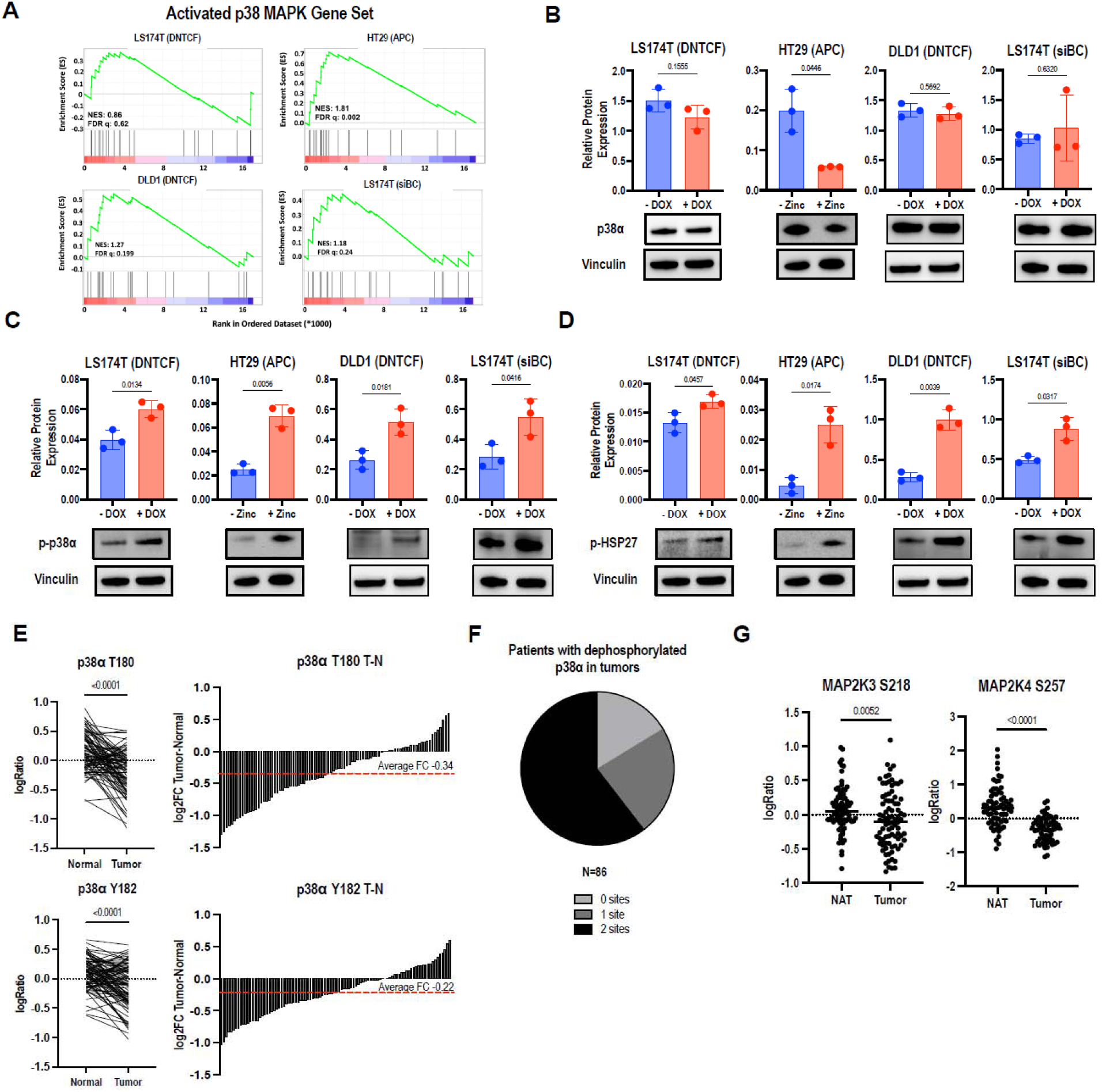
p38 signaling is inhibited by APC/β-catenin/TCF signaling. (A) GSEA analysis to identify p38-activated genes revealed an enrichment of p38 activity in the absence of APC/β-catenin/TCF signaling in the four CRC conditional cell lines. (B) p38α expression was unchanged in LS174T (DNTCF), LS174T (siBC), and DLD1 (DNTCF) cells, and was reduced in HT29 (APC) cells under +DOX/+Zinc conditions in CRC conditional cell lines. (C) p-p38α expression was increased in +DOX/+Zinc conditions of CRC conditional cell lines. (D) p-p38α downstream activity marker p-HSP27 was increased in +DOX/+Zinc conditions of CRC conditional cell lines. (E) Phosphorylation of p38α sites threonine (T)180 and tyrosine (Y)182 are reduced in CRC tumors compared to matched normal colon samples in CPTAC phosphoproteomics. (F) 52 patients (61%) exhibit reduced phosphorylation at both p38α phosphosites, while 20 patients (23%) exhibit reduced phosphorylation at one site, and 14 patients (16%) have no change or increases at both phosphosites in tumors relative to matched normal tissue. (G) Phosphorylation of protein kinases MKK3 and MKK4 at phosphosites S218 and S257 are reduced in CRC compared to matched normal colon samples in CPTAC. Statistics were calculated using an unpaired Welch’s *t* test (B-D, G) or a paired Student’s *t* test (E). Error bars represent the standard deviation (SD).

### The p38α signaling network is inhibited in human colorectal tumors

Activation of p38α occurs via dual phosphorylation of the Thr-Gly-Tyr motif in the activation loop by mitogen activated protein kinase kinase 3 (MKK3), MKK4, and MKK6.^39^ Although p38α mRNA (TCGA) and total protein expression are unchanged in colorectal tumors, compared to matched normal tissue (Supplemental Fig 4A, B), phosphoproteomic analysis from the Clinical Proteomic Tumor Analysis Consortium (CPTAC)^40^ revealed a reduction of p38α phosphorylation at both T180 (n=86) and Y182 sites (n=96) in tumors (Fig. 3E). Among these tumors, 61% exhibited reduced phosphorylation at both sites, 23% exhibit reduced phosphorylation at one site and 16% were without reductions (Fig. 3F). Consistent with these findings, phosphorylation of upstream kinases MKK3 and MKK4 at phosphosites S218 and S257, respectively, associated with protein activation, was reduced in colorectal tumors compared to normal mucosa (Fig. 3G).^41,42^ Thus, mutant APC/β-catenin/TCF signaling, which drives transformation in most CRC, inhibits p38α phosphorylation and activity in cells and tumors. In turn, inhibition of p38α contributes to loss of GUCA2A and GUCA2B, functionally silencing the GUCY2C tumor-suppressor receptor during early tumorigenesis.

### Beyond GUCA2A/B, most APC/β-catenin/TCF-regulated gene expression requires p38α

Transcriptomic analysis in LS174T (siBC), LS174T (DNTCF), and HT29 (APC) cells revealed a substantial subset of genes regulated by APC/β-catenin/TCF signaling which was mediated by p38α (Fig. 4A, Supplemental Fig. 5A, C). In wildtype (WT) p38α cells, pathway analysis revealed that silencing APC/β-catenin/TCF enriched TNFα, KRAS, and estrogen signaling (Fig. 4B, Supplemental Fig. 5B, D). Conversely, activating APC/β-catenin/TCF enriched pathways such as MYC, APC/β-catenin, and E2F (Fig. 4B, Supplemental Fig. 5B, D). Of the 115 genes consistently regulated by APC/β-catenin/TCF signaling across the three p38α-WT models, 93 (80%) were no longer significantly regulated following p38α deletion (Fig. 4C, D). These 93 genes constituted the p38α-dependent APC/β-catenin/TCF transcriptional signature. Analysis of Motif Enrichment (AME)^43^ revealed that this set of jointly regulated genes shared motifs for transcription factors controlled by p38α in their 5’ regulatory regions (Fig. 5A, B).^44–46^ p38α-regulated transcription factors SP3 and KLF16 were among the top three motifs enriched uniquely in the p38α regulated gene set, shared by 72% and 67%, respectively, of genes in the set (Fig. 5A, B). Further, ARID5A, SP family and KLF family transcription factors operate downstream of p38α, and their consensus motifs were enriched across this gene set (Fig. 5B).^47–49^ Indeed, 99% of genes in this set contained at least one of the top ten enriched consensus motifs in their promoters, and 73% contained at least four (Fig. 5C). The relevance of this p38α-dependent APC/β-catenin/TCF transcriptomic signature is highlighted by analysis of the colorectal adenocarcinoma cohort from The Cancer Genome Atlas (TCGA-COAD), which revealed that 59% of this gene set (53 genes) was differentially expressed in tumors compared to normal colorectum (Fig. 5D, E). Within the gene signature enriched in TCGA-COAD, 23 genes were upregulated and 30, including GUCA2A, were downregulated in tumors (Fig. 5D). Finally, GSEA against TCGA-COAD revealed that overall, the p38α-dependent gene signature was enriched in healthy tissue relative to tumors, indicating repression of p38α-regulated genes in CRC (Fig. 5F). Together, these observations suggest that p38α mediates this APC/β-catenin/TCF-regulated transcriptional program through p38-responsive transcription factors, including SP- and KLF-family proteins associated with transcriptional complexes at target promoters and enhancers.

**Figure 4.**
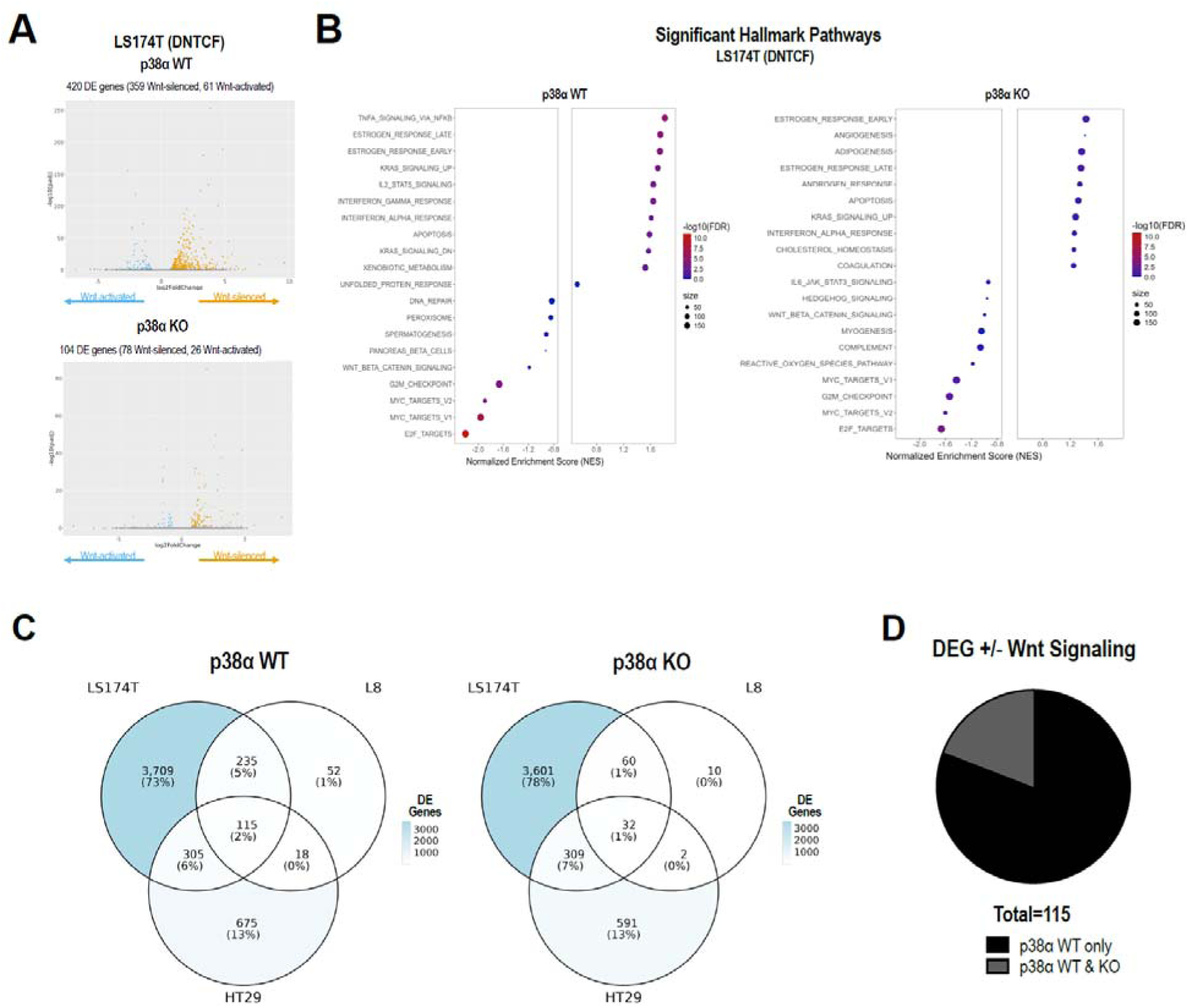
p38α is necessary for regulation of APC/β-catenin/TCF-dependent gene expression in CRC cells. (A) Volcano plots of genes silenced or activated by APC/β-catenin/TCF signaling in LS174T (DNTCF) cells with or without p38α. In p38α WT cells, 420 genes are differentially expressed (359 Wnt-silenced, 61 Wnt-activated) while in p38α KO cells, 104 genes are differentially expressed (78 Wnt-silenced, 26 Wnt-activated). (B) Significant hallmark pathways in LS174T (DNTCF) p38α WT and p38α KO cells. (C-D) Differentially expressed genes in LS174T (DNTCF), LS174T (siBC), and HT29 (APC) cells. In p38α WT cells, 115 shared genes are differentially expressed, while in p38α KO cells, 32 shared genes are differentially expressed. Of the 32, 22 overlap with the p38α WT group (10 genes differentially expressed in p38α KO cells only).

**Figure 5.**
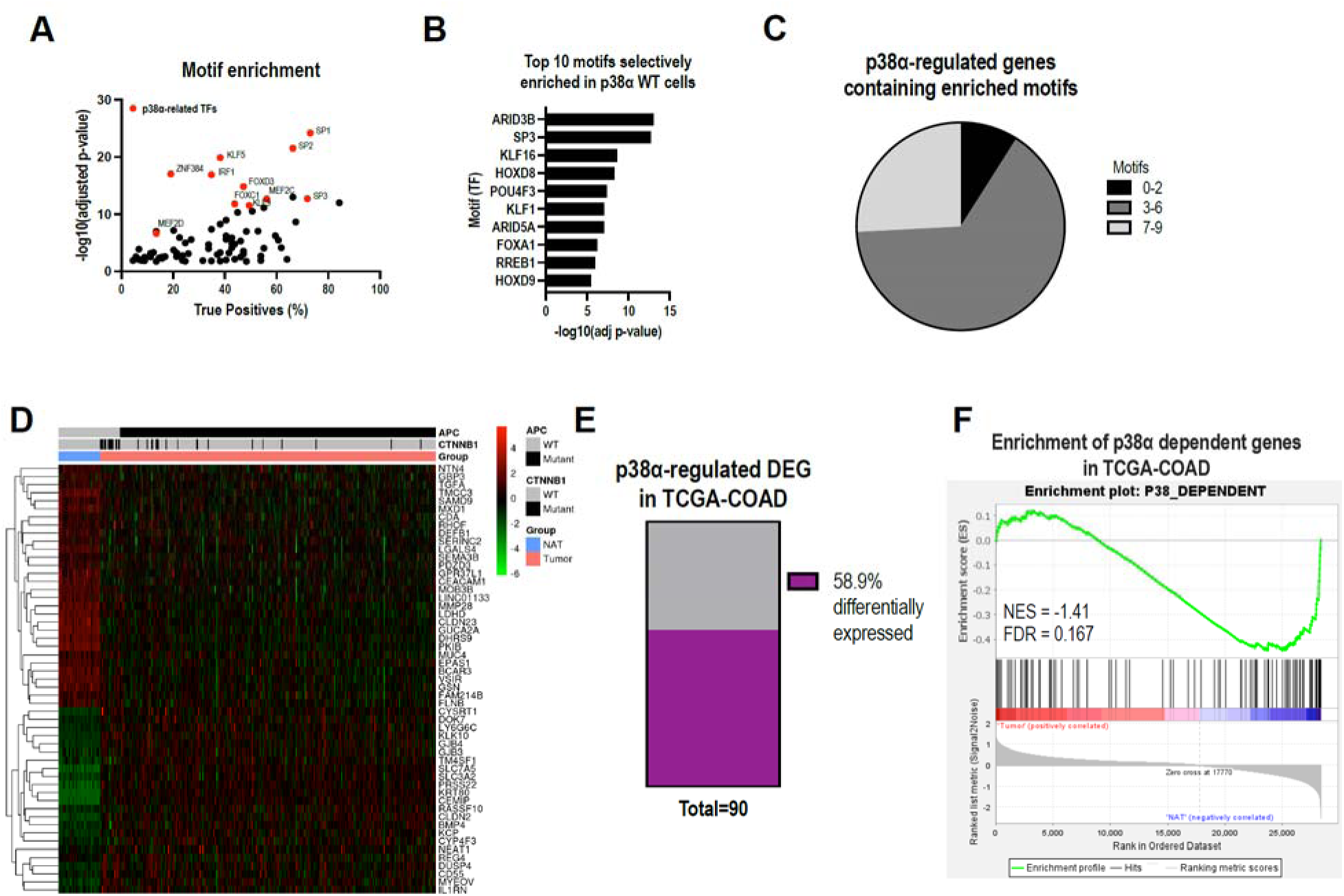
The p38α-dependent gene signature is enriched in TCGA-COAD. (A) AME analysis identified enriched transcription factor binding motifs in the 1000 bp putative promoters of p38α-dependent, APC/β-catenin/TCF-regulated genes. Transcription factors associated with p38α signaling are identified in red. (B) The top 10 enriched motifs unique to the p38α-dependent gene signature. (C) Distribution of the top 10 enriched motifs across genes in the p38α dependent gene signature. Of these genes, 9% contain 0-2 motifs, 65% contain 3-6 motifs, and 26% contain 7-9 motifs within their putative promoters. (D) Enrichment of the p38α-dependent gene signature in the TCGA COAD cohort. (E) 58.9% of the gene signature was differentially expressed between NAT and tumor tissue in TCGA COAD. (F) GSEA of the gene signature demonstrates enrichment in NAT compared to tumor tissue in TCGA COAD (NES = - 1.41, FDR=0.167).

## Discussion

The GUCY2C hormones GUCA2A and GUCA2B are universally lost in colorectal cancer.^2,21,23,24^ An LCR in the *GUCA2A* upstream regulatory region coordinates hormone transcription and is regulated by APC/β-catenin/TCF signaling.^2^ Here, we reveal that p38α is necessary for GUCA2A and GUCA2B expression, using pharmacological inhibition, genetic ablation, and rescue of p38α expression (Fig. 1, 2). APC/β-catenin/TCF signaling reduces p38α phosphorylation and activity and is associated with reduced enrichment of p38α at the GUCA2A LCR, which correlates with loss of GUCY2C hormones in CRCs. In that context, phosphorylation of p38α and its upstream kinases MKK3 and MKK4 is reduced in CRC compared to normal colonic mucosa (Fig. 3). Transcriptomic profiling identified a subset of APC/β-catenin/TCF-dependent genes that require p38α expression, revealing that this kinase regulates a substantial component of that oncogenic transcriptional program driving CRC (Fig. 4). Promoter analysis of this gene set revealed enrichment of shared transcription factor motifs, including those regulated by p38α (Fig. 4). These observations define p38α as a critical node at the intersection of reciprocal regulation of APC/β-catenin/TCF and GUCY2C axes. Further, they establish a previously unrecognized link between p38 signaling and canonical oncogenic gene programs driving transformation in CRCs.

Although p38α shares substantial sequence homology with other p38 isoforms (∼75% with p38β), accumulated evidence indicates that p38 signaling can be both tissue-dependent and isoform-specific.^50–55^ Indeed, distinct roles among p38 isoforms have been defined in different models, including opposing roles for p38α and p38γ in the regulation of c-Jun activity.^56^ Further, in contrast to our findings of p38α, p38γ is required for APC/β-catenin/TCF signaling, promoting inflammation-associated colorectal tumorigenesis.^57^ Here, regulation of a substantial subset of genes by APC/β-catenin/TCF signaling is specifically mediated by p38α compared to other p38 isoforms. In that context, while other isoforms of p38 are expressed in p38α KO cells, they do not compensate for the loss of p38α in regulating gene expression by APC/β-catenin/TCF signaling. Indeed, expression of GUCA2A and GUCA2B is lost despite signaling from other p38 isoforms (Fig. 2). Additionally, eliminating the other p38 isoforms together with p38α does not further impact expression of GUCA2A or GUCA2B (Fig. 2). Our results support a unique, non-redundant role for p38α in controlling the core signature of APC/β-catenin/TCF-regulated genes in CRC cells.

The present study defines direct negative crosstalk between p38α and Wnt/APC/β-catenin/TCF signaling in CRC cells. In contrast, previous reports have described p38 as a positive regulator of APC/β-catenin/TCF signaling. In totipotent mouse F9 teratocarcinoma cells, Wnt3a binds to Frizzled and activates Gα_q_ and Disheveled, promoting transient activation of p38.^58^ In mesenchymal cells, Wnt3a and Wnt4 activate p38 to promote osteogenic differentiation.^59^ In addition, p38 phosphorylates GSK3β, inhibiting its kinase activity and promoting β-catenin nuclear translocation.^60–62^ Further, p38α directly phosphorylates nuclear β-catenin and facilitates its binding to Wnt target gene promoters in colorectal cancer cells.^63^ In contrast to earlier studies, we find that silencing p38α promotes an APC/β-catenin/TCF transcriptional program in three CRC cell lines (Fig. 4). Further, there is a complex interaction between APC/β-catenin/TCF and p38α signaling pathways in CRC. Specifically, APC/β-catenin/TCF signaling suppresses p38α phosphorylation and activation, while GUCA2A and GUCA2B, downstream of APC/β-catenin/TCF signaling, require p38α activity for gene expression. Consistent with this, hormone activation of their cognate receptor GUCY2C stimulates p38α signaling and induces cellular senescence.^64^ Together, these observations place p38α at the intersection of APC/β-catenin/TCF signaling and the GUCY2C tumor-suppressor axis and support a broader reciprocal relationship between these oncogenic and homeostatic pathways. We propose that p38α regulation occurs downstream of TCF, as APC/β-catenin-driven gene expression in these cells depends on TCF activity. Future studies will determine whether APC/β-catenin/TCF signaling regulates p38α through transcriptional control of upstream p38α pathway components, modulation of growth factor signaling, or other intermediates that converge on the p38α cascade.

Moreover, transcriptomic profiling supports an expansion of this model, revealing that p38α is necessary for regulation of a substantial component of the shared APC/β-catenin/TCF transcriptional signature defined across these CRC models. Notably, more than half of these p38α- and APC/β-catenin/TCF-dependent genes are differentially expressed in colorectal tumors compared to normal tissue in TCGA datasets, underscoring their relevance in human CRC. Promoter analysis of this gene set revealed enrichment of motifs including p38α-regulated transcription factors SP1, KLF5, MEFs, and AP-1, suggesting that this kinase regulates oncogenic APC/β-catenin/TCF-driven gene expression through a convergent transcription factor network. Future studies will define the transcription factors mediating p38α-dependent APC/β-catenin/TCF target gene regulation.

These observations raise important questions regarding how p38 signaling contributes to colorectal tumorigenesis, where it has been associated with both tumor suppression and tumor promotion. Interestingly, p38α has been implicated as a tumor suppressor in multiple models of colon tumorigenesis.^10,65^ Beyond CRC, loss of upstream p38α kinases MKK3 and MKK6 amplifies tumor growth in nude mice.^66^ Conversely, p38α, p38γ, and p38δ are pro-tumorigenic in inflammation-associated CRC.^57,65,67^ Additionally, a p38 inhibitor reduced tumor growth in a xenograft nude mouse model and a 4 month-old azoxymethane-treated Apc^Min^ model.^68,69^ In the context of the present observations, these models using p38 inhibitors with established tumors suggest opposing roles for p38α in tumor initiation and progression. Opposing roles for p38α, protective against CRC initiation but oncogenic in tumor promotion, have been suggested.^65^

Finally, these results underscore the convergent roles for p38α and GUCY2C axes in normal intestine. p38α is expressed in a gradient along the crypt-surface axis, with maximal expression in differentiated epithelial cells near the surface, whereas the APC/β-catenin/TCF signaling gradient is maximal in the undifferentiated crypt compartments.^1,7^ Further, GUCA2A and GUCA2B also are expressed in a gradient along the crypt-surface axis, closely matching that of p38α.^23^ Here, activating the Wnt pathway in normal colonoids reduced p38α signaling and suppressed GUCY2C hormone expression (see Fig. 1G-H). In that context, GUCY2C is a tumor suppressive signaling axis, and loss of its cognate hormones drives colorectal transformation.^16–18^ Here, we identify a mechanism by which a key initiating transforming event (e.g., mutations in APC) in colorectal tumorigenesis simultaneously activates oncogenic β-catenin/TCF programs and disables a homeostatic tumor-suppressor pathway. APC/β-catenin/TCF-mediated inhibition of p38α suppresses GUCA2A and GUCA2B transcription, functionally silencing GUCY2C despite preservation of its receptor. Thus, loss of p38α signaling provides a mechanistic link between APC pathway activation and extinction of the GUCY2C tumor-suppressor axis and reinforces reconstitution of hormone expression as a potential strategy to restore this homeostatic circuit to oppose early colorectal tumorigenesis.

## Materials and Methods

### Cell and Organoid Culture Reagents

McCoy’s 5A (#10050CV), Dulbecco’s Modified Eagle Medium (DMEM) (#10013CV), Advanced DMEM/F12 (#), Matrigel (#354230), and cell recovery solution (#354253) were from Corning (Corning, NY). Ca2+/Mg2+-free Hank’s Balanced Salt Solution (HBSS) (#14175095), GlutaMax supplement (#35050061), penicillin-streptomycin (#15140122), HEPES (#15630080), HyClone fetal bovine serum (#SH3007103), hygromycin B (#10687010), zeocin (#R25005), blasticidin hydrochloride (#A1113903), geneticin/G418 sulfate (#10131027), doxycycline hydrochloride (#BP2653-1), puromycin (#A1113803), N-2 supplement (#17502048), and B-27 supplement (#12587010) were from Thermo Fisher Scientific (Waltham, MA). N-acetyl-l-cysteine (#A9165), A83-01 (#SML0788), nicotinamide (#NO636), and zinc chloride (#Z0152) were from Sigma-Aldrich (St Louis, MO). Wnt3a-, R-Spondin-, and Noggin-conditioned medium (from L-WRN cells; #CRL-3276) was from the American Type Culture Collection (Manassas, VA). Mouse recombinant epidermal growth factor (EGF) (#315-09) and Noggin (#250-38) were from PeproTech (Rocky Hill, NJ). Human recombinant R-spondin-1 (#4645-RS-025/CF) and SB202190 (#1264) were from R&D Systems (Minneapolis, MN). CHIR99021 (#129830-38-2) and Y-27632 (#252917-06-9) were from StemCell Technologies (Vancouver, Canada). SB203580 (#13067) and trametinib (#16292) were from Cayman Chemical Company (Ann Arbor, MI). Single guide RNAs (sgRNAs) and Cas9 protein for CRISPR/Cas9 gene editing were from Synthego (Redwood City, CA). pCDNA3 Flag p38 alpha was a gift from Roger Davis (Addgene plasmid #20351; http://n2t.net/addgene:20351; RRID:Addgene_20351).

### Human Organoid Studies

De-identified human colon biopsies were obtained from the Jefferson Biobank and washed in cold HBSS supplemented with 0.5% penicillin-streptomycin. The colonic mucosa was dissected from submucosa and minced into ∼1 cm pieces. It was then incubated shaking in HBSS supplemented with 20 mM EDTA at 4°C for 30 minutes. It was then vortexed for 2 min, letting incubate for 5min on ice after each minute. Crypts were spun down at 125 *g* for 3 min, and then washed in sorbitol twice, spinning down at 125 *g* for 3 min at 4°C after each wash. Finally, crypts were washed in Advanced DMEM/F12 and plated in 50 μL of Matrigel in 24-well plates. Matrigel patties were incubated for 15-20 min at 37°C and then overlaid with 500 μL of complete medium supplemented with Wnt3a-, R-spondin-, and Noggin-conditioned medium (L-WRN, 1:1 dilution), 1× N-2, 1× B-27, 100 μM N-acetyl-l-cysteine, 50 ng/mL EGF, 500 nM A83-01, 10 μM CHIR99021, and 10 μM Y-27632. CHIR99021 and Y-27632 were removed from medium 48 h after passaging. Organoids were passaged weekly. For organoid differentiation studies, medium was replaced with differentiation medium (complete medium supplemented with 1 μg/mL R-spondin-1, 100 ng/mL Noggin, 1× N-2, 1× B-27, 100 μM N-acetyl-l-cysteine, 50 ng/mL EGF, 500 nM A83-01, and 10 μM Y-27632). Differentiation medium was replaced daily for 3 days. To harvest, organoids were lysed in M-PER or RLT buffer for protein or RNA analyses, respectively.

### Cell Lines

DLD1 and LS174T cells stably expressing Tet-inducible dominant negative TCF7L2 (DNTCF), and LS174T cells stably expressing Tet-inducible shRNA to β-catenin were provided by Dr. H. Clevers in November 2013. Cells were cultured in DMEM supplemented with 10% fetal bovine serum (FBS), 500 μg/mL zeocin, and 10 μg/mL blasticidin. Cells were induced with 1 μg/mL DOX for 24 hours (DLD1-DNTCF, LS174T-DNTCF), or 72 h (LS174T-siBC). HT29 cells stably expressing zinc-inducible wild-type APC were provided by Dr. B. Vogelstein in September 2013. HT29 cells were cultured in McCoy’s-5A supplemented with 10% FBS and 600 μg/mL hygromycin. Cells were induced with 300 μM zinc chloride for 24 h. Conditional cell lines were authenticated at each use by testing their genetic inducibility, and all cell lines were confirmed to be free of mycoplasma semiannually.

### Immunoblots

Protein was extracted from total cell lysates using M-PER (Thermo Fisher Scientific; #78501) supplemented with protease and phosphatase inhibitors (Thermo Fisher Scientific; #A32955 or #78440). Lysates were prepared in 4X LDS Sample buffer (Invitrogen, Waltham, MA; #NP0007) supplemented with 2.75 mM 2-Mercaptoethanol (Thermo Fisher Scientific; #21985023), heated at 95°C for 10 min, loaded onto 4%– 12% Bis-Tris gels (20 μg/lane; Thermo Fisher Scientific; #NP0336), and transferred to nitrocellulose membranes with an iBlot2 gel transfer device (Thermo Fisher Scientific; IB21001). Blots were blocked for 1 h in tris-buffered saline (TBS) containing 0.1% Tween 20 and 5% BSA, then incubated overnight at 4°C with primary antibodies listed in Supplemental Table 1. Secondary antibodies conjugated to horseradish peroxidase were used at 16 pg/μL (Jackson Immuno Research Laboratories, West Grove, PA; #111-035-144 or #115-035-062) or 0.32 ng/μL (Jackson Immuno Research Laboratories, West Grove, PA; #016-030-084). Blots were developed with Dura or Femto Chemiluminescent Substrates (Thermo Fisher Scientific; #34075, #34095) and imaged on a ChemiDoc MP Imaging Station (Bio-Rad, Hercules, California). Bands were quantified by densitometry normalized to that of HSP90 or vinculin using Bio-Rad imaging software.

### RNA Analysis

Samples were lysed in RLT buffer supplemented with 550 μM 2-mercaptoethanol and RNA was purified on spin columns using the RNeasy kit (Qiagen, Hilden, Germany; #74104) according to kit directions. RNA concentration and purity were measured with a Nanodrop 1000 (Thermo Fisher Scientific), and 200 ng of RNA was reverse transcribed to complementary DNA (cDNA) using the TaqMan Reverse Transcription kit according to kit directions (Thermo Fisher Scientific; #N8080234). Transcripts were quantified by qRTPCR using TaqMan primer probes on a QuantStudio™ 3 Real-Time PCR System (Thermo Fisher Scientific), with TaqMan Universal PCR Master Mix (Thermo Fisher Scientific; #4318157), per kit instructions. Primer probes are listed in Supplemental Table 1.

### RNA Sequencing

Total RNA was extracted as above, and library preparation and sequencing were performed by Azenta Life Sciences. The RNA sample received was quantified using Qubit 4.0 Fluorometer (ThermoFisher Scientific, Waltham, MA, USA) and RNA integrity was checked using TapeStation (Agilent Technologies, Palo Alto, CA, USA). The NEBNext Ultra II RNA Library Prep Kit for Illumina & NEBNext Poly(A) mRNA Magnetic Isolation Module (New England Biolabs, Ipswich, MA, USA), including clustering and sequencing reagents, was utilized according to the manufacturer’s recommendations. Briefly, mRNAs were initially enriched with Oligod(T) beads. Enriched mRNAs were fragmented for 15 minutes at 94°C. First strand and second strand cDNA were subsequently synthesized. cDNA fragments were end repaired and adenylated at 3’ ends, and universal adapters were ligated to cDNA fragments, followed by index addition and library enrichment by PCR with limited cycles. The sequencing library was validated on the Agilent TapeStation (Agilent Technologies, Palo Alto, CA, USA) and quantified by using Qubit 4.0 Fluorometer (ThermoFisher Scientific, Waltham, MA, USA) as well as by quantitative PCR (KAPA Biosystems, Wilmington, MA, USA). The sequencing libraries were multiplexed and clustered onto a flowcell on the Illumina NovaSeq X Plus instrument according to manufacturer’s instructions. The samples were sequenced using a 2×150bp Paired End (PE) configuration. Image analysis and base calling were conducted by the NovaSeq Control Software (NCS). Raw sequence data (.bcl files) generated from Illumina NovaSeq was converted into fastq files and de-multiplexed using Illumina bcl2fastq 2.20 software. One mismatch was allowed for index sequence identification.

Trimming, alignment, and expression quantification were performed with in-house Python scripts implemented with Snakemake.^70^ Raw reads were trimmed with Trimmomatic^71^ to remove low-quality bases. Clean reads were aligned to the human (GRCh38) reference genome using STAR (v2.7.4a)^72^ and reads mapping to each gene were counted with RSEM (v1.3.3)^73^ in un-stranded mode. Differential gene expression (adjusted p value <.05 and log fold-change threshold = log_2_(1.5)) was determined between cells treated with or without doxycycline (LS174T and L8) or ZnCl (HT29) and between cells with or without p38α, using the DESeq2 (v1.42.1)^74^ package in R (R Foundation for Statistical Computing, Vienna, Austria, v4.3.1). The default normalization of median-of-ratios was applied with DESeq2 before differential expression testing. Heat maps were generated using pheatmap (v1.0.13). Gene set enrichment analysis (GSEA)^75^ was performed for each cell line, comparing genes (ranked by log-fold change) to the H:Hallmark Gene Sets in MSigDB (v7.5.1)^76^ with fgsea (v1.28.0)^77^, and results were plotted using the ggplot2 (v4.0.2) package in R. GSEA performed with the p38α-dependent gene set against TCGA-COAD data was run in phenotype mode with statistical significance evaluated as FDR<0.25. 3 genes from the p38α gene signature (*GRAMD1B*, *KRT7*, *MUCL3*) were absent from the TCGA-COAD transcriptomics and thus excluded from the analysis. TCGA tumor samples with no APC pathway mutations (*APC*, *CTNNB1*, *RNF43*, *ZNRF3*, *AXIN1*, *AXIN2*)^78^ were not included in the analysis. RNA sequencing data have been deposited in the Gene Expression Omnibus (GSE348607) and will be made publicly available upon publication of this manuscript.

### CRISPR/Cas9 Gene Editing

CRISPR/Cas9-mediated gene knockout in colorectal cancer cell lines was performed using the Lonza Nucleofector system. Target sequences for the desired gene knockout were designed by Synthego (Redwood City, CA) and are listed in Supplemental Table 1. CRC cell lines were cultured in appropriate growth medium until they reached approximately 70-80% confluency. Prior to nucleofection, cells were harvested by trypsinization, counted, and resuspended in Nucleofector solution. For each nucleofection, 250,000 cells were mixed with spCas9-2NLS (Synthego) and sgRNA. The mixture was transferred into a nucleocuvette and nucleofected using the optimized program on the Lonza 4D-Nucleofector System. After nucleofection, cells were transferred into pre-warmed RPMI/10% FBS medium and incubated overnight at 37°C with 5% CO_2_. The next day, medium was replaced with normal growth medium. Single-cell clones were isolated by limiting dilution. Genomic DNA from polyclonal populations and individual clones was extracted using a genomic DNA extraction kit (Invitrogen Purelink Genomic DNA Mini Kit), and gene knockout was confirmed by PCR amplification and Sanger sequencing of the target locus. Additionally, Western blot analysis was performed to verify the absence of the target protein.

### Kinase-Null and Phosphomimetic Constructs

The p38α sequence from pCDNA3 Flag p38 alpha was cloned into the pCDH-EF1α-MCS third-generation lentiviral backbone (System Biosciences #CD502A-1) by GenScript Biotech.^35^ Kinase-null and phosphomimetic mutations were designed based on Diskin et al: the kinase-null construct contains the mutations T180A and Y182F, and the constitutively active phosphomimetic construct contains the mutations D176A and F327S. Plasmids were packaged into lentivirus and transduced into LS174T (DNTCF) cells as described previously.^36,79^

### Generating the BirA/Cas9 Cells

DLD1 (DNTCF) cells were transfected with pEF1a-BirA-V5-neo (Addgene #100548) and pEF1a-FB-dCas9-puro (Addgene #100547) using FuGENE transfection reagent according to the manufacturer’s instructions (Promega E2311). Cells were selected and maintained with 1 µg/mL puromycin and 600 µg/mL Geneticin. For each experiment, stably expressing birA/FBdCas9 cells were transduced with 3^rd^ generation lentivirus containing a gRNA targeting the GUCA2A promoter (sequence: GCCCGGCTGTGGGTCAAAGTAGG, 167-190bp upstream of the transcription start site)^2^ or the non-mammalian GAL4 (Addgene #100549) as described previously^75^, and treated with or without 1 µg/mL doxycycline for 24 h to induce DNTCF.

### Chromatin Immunoprecipitation (ChIP)-qPCR

To validate FBdCas9 binding at the targeted locus, chromatin immunoprecipitation was performed as previously described.^2^ For each IP, 25 µL Dynabeads MyOne Streptavidin T1 (Invitrogen #65602) were incubated with 25 µg chromatin while rotating overnight at 4°C. Following DNA isolation and purification, 2 µL post-IP or input DNA and 200 nM forward and reverse primers were used per qPCR reaction using PowerUp SYBR Green Master Mix (Applied Biosystems #A25776) according to the manufacturer’s instructions in a QuantStudio 3 Real-Time PCR System (Applied Biosystems). Quantification was performed using the percentage of input method.

### CAPTURE-Immunoblot

Streptavidin immunoprecipitation was performed following the CAPTURE-Proteomics protocol^37^ with the following modifications: ∼40×10^6^ cells were crosslinked in 2% formaldehyde for 10 min (Thermo Scientific #28908), then quenched in 0.25 M glycine for 5 min. Cells were washed twice with cold DPBS, then lysed in 10 mL Cell Lysis Buffer. Lysates were rotated for 15 min at 4°C, then centrifuged at 2,300g for 5 min at 4°C to isolate nuclei. Nuclei were resuspended in 2.5 mL Nuclear Lysis Buffer and incubated for 10 min at room temperature. The suspension was mixed with 7.5 mL 8M urea and centrifuged at 16,100g for 25 min at room temperature. The pellet was again resuspended in 2.5 mL Nuclear Lysis Buffer, mixed with 7.5 mL of 8 M urea, and centrifuged, a total of 4 times. The pellet was finally washed with 2.5 mL Cell Lysis Buffer, centrifuged again at 16,100g for 25 min at room temperature, and resuspended in 1 mL IP Binding Buffer with no NaCl. Nuclei were sonicated to an average length of 500 bp at 4°C (QSonica Q800R2). Chromatin was then combined and centrifuged at 16,100g for 25 min at 4°C. NaCl was added to a final concentration of 150 mM. For each IP, 150 μL of streptavidin agarose slurry (Invitrogen #SA10004) was washed 3 times in 1 mL of IP Binding Buffer, then incubated with 0.33 mg chromatin at 4°C overnight in a rotator. Streptavidin beads were then collected by centrifugation at 800g for 3 min at 4°C. Beads were washed 5 times with IP Binding Buffer containing NaCl and 5 times with 50 mM Tris. Following the washes, beads were resuspended in 70 μL XT sample buffer (BioRad #1610791) containing 1.25% β-mercaptoethanol, boiled for 20 minutes, and loaded on an SDS-PAGE gel for analysis by immunoblot.

### Motif Analysis

Promoter motif enrichment analysis was performed using the MEME Suite.^43^ Briefly, 1000 bp putative promoter sequences upstream of the transcription start site (TSS) for p38α-dependent and independent genes of interest were batch extracted from the hg38 reference genome using the UCSC Genome Browser. 89 and 31 promoter sequences were successfully extracted, respectively, and provided as input for the Analysis of Motif Enrichment (AME) tool within MEME Suite. AME was used to identify significantly enriched transcription factor binding motifs in the promoter sequences relative to shuffled input sequences using the 2018 JASPAR non-redundant vertebrate DNA motif database.^43,80,81^ Motif enrichment was scored using the average odds score method, in which sequences scoring above a motif-specific threshold were classified as true positives. Enrichment was assessed by comparing the proportion of true-positive sequences between the promoter and shuffled control sequence sets using Fisher’s exact test, with Bonferroni-adjusted p-values reported to account for multiple hypothesis testing. The reported list of top ten enriched motifs consists of those which did not overlap with enriched motifs in the p38α-independent signature.

### siRNA Knockdown

Cells were transfected with siRNA using Lipofectamine RNAiMAX Transfection Reagent (Invitrogen #13778075) according to the manufacturer’s instructions. Briefly, cells were seeded 24 hours prior to transfection to achieve approximately 50-70% confluency at the time of transfection. siRNA duplexes and Lipofectamine RNAiMAX reagent were separately diluted in Opti-MEM reduced serum medium and incubated for 5 min at room temperature prior to combining. The siRNA-lipid complexes were incubated for 10-20 min at room temperature and then added dropwise to cells. Cells were incubated for 48-72 h prior to downstream analyses. A non-targeting siRNA control was included in all experiments, and knockdown efficiency was confirmed by qRT-PCR. siRNAs are listed in Supplemental Table 1.

### Luciferase Reporter Assay

Luciferase reporter assays were performed as previously described by Rappaport et al.^2^ Briefly, cells were seeded in white 96-well plates and cotransfected with Firefly and Renilla luciferase reporter plasmids (pRLTK; Promega; #E2241), using Fugene HD (Promega; #E2311), according to kit instructions. DOX was added to induce L8 (DNTCF) cells. A total of 48 h after transfection, cells were lysed with the Dual-Glo Luciferase Assay kit (Promega; #E2940), following kit instructions. Luminescence was read on a FLUOstar Omega plate reader (BMG Labtech). Firefly luciferase activity was normalized to Renilla luciferase activity in each well.

### Statistics

Results are presented as the mean ± the standard deviation (SD), and a p-value of <0.05 was considered significant. Statistical analysis was performed in GraphPad Prism 11 (Version 11.0.0) or in R. Data were analyzed via one-way and two-way ANOVAs with Tukey’s multiple comparisons tests, and two-tailed unpaired Welch’s t tests.

## Acknowledgements

The results published here are in part based upon data generated by The Cancer Genome Atlas Research Network (https://www.cancer.gov/tcga), the Clinical Proteomic Tumor Analysis Consortium (https://dctd.cancer.gov/research/networks/cptac), and the National Center for Biotechnology Information Gene Expression Omnibus (https://www.ncbi.nlm.nih.gov/geo). LS174T (siβ-catenin), LS174T (DNTCF), and DLD1 (DNTCF) cell lines were provided by H. Clevers. HT29 (APC) cells were provided by B. Vogelstein. S.A.W. was supported by funding including from the NIH (1R01 CA204881, 1R01 CA206026, 1R21 1NS130388, 1R01 DK1388341), DoD (W81XWH-17-PRCRP-TTSA), the American Parkinson Disease Association, Parkinson’s Foundation Impact Award PF-IMP-1045175, and Targeted Diagnostic and Therapeutics, Inc. A.E.S. was supported by funding including from the DoD (W81XWH-19-1-0263, W81XWH-22-1-0207), NIH (1R21 CA267087, 1R21 CA286339), and the Kleberg Foundation. S.A.W. and A.E.S. were additionally supported by funding from The Courtney Ann Diacont Memorial Foundation and Lorraine and David Swoyer. A.A.E, A.E.E, and A.S.D. were supported by the T32 institutional award for Postdoctoral Training in Clinical Pharmacology (GM008562) and A.A.E was additionally supported by the institutional Training Program in Alcohol Research (AA007463). A.C. was supported by the institutional Training Program in Cancer Biology (T32 CA236736). T.J.M.K. was supported by the PhRMA Foundation Predoctoral Fellowship in Drug Discovery. A.E.S. and data utilizing the Biostatistics, Bioinformatics, Research Informatics Shared Resource at the Sidney Kimmel Comprehensive Cancer Center at Jefferson Health were supported by the National Cancer Institute of the National Institutes of Health under Award Number P30 CA056036. The content is solely the responsibility of the authors and does not necessarily represent the official views of the NIH.

## Supplemental Figures

**Supplemental Figure 1.**
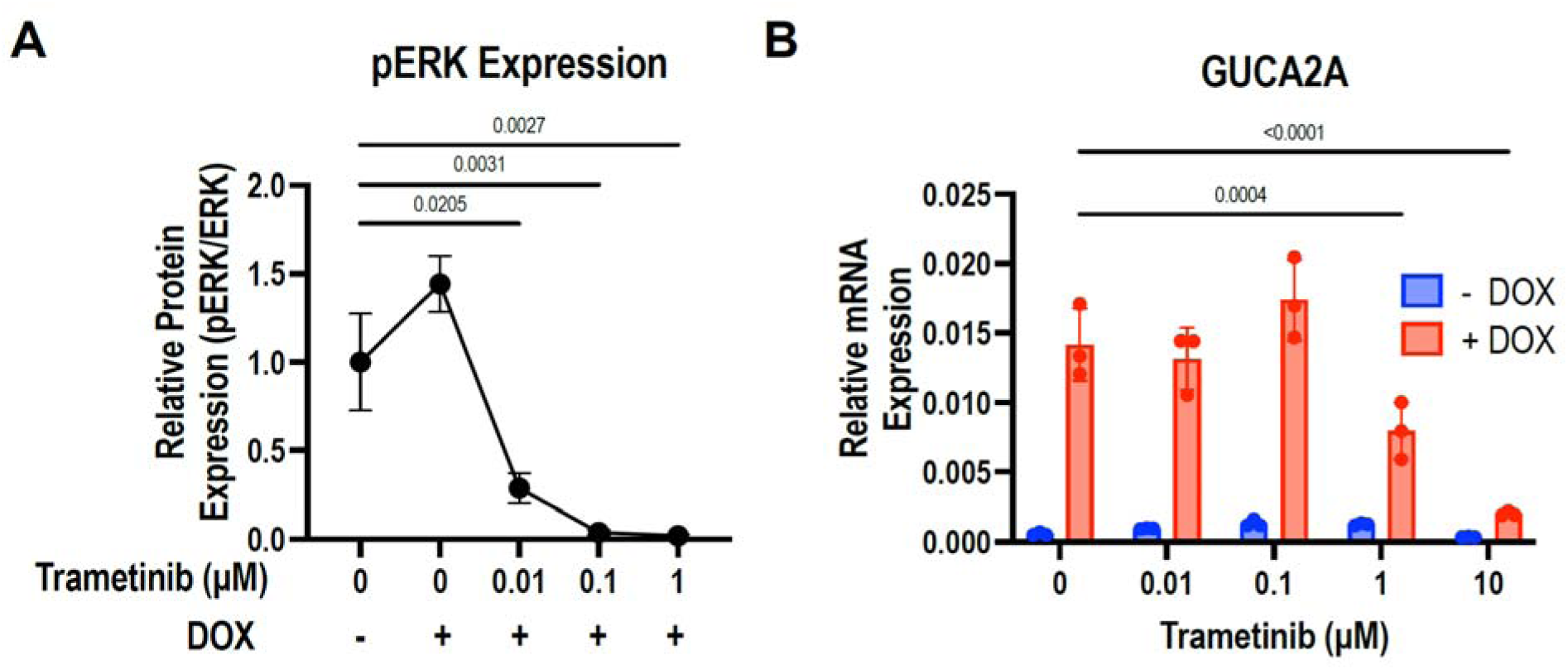
ERK signaling is not required for GUCY2C hormone expression. (A) Phosphorylation of ERK1/2, measured by normalizing phosphorylated ERK to total ERK, was inhibited with increasing concentrations of the MEK1/2 inhibitor trametinib. (B) GUCA2A mRNA expression was inhibited at higher concentrations of trametinib (1 mM and 10 mM). Statistics were calculated using a two-way ANOVA with Tukey’s multiple comparisons test. Error bars represent the standard deviation (SD).

**Supplemental Figure 2.**
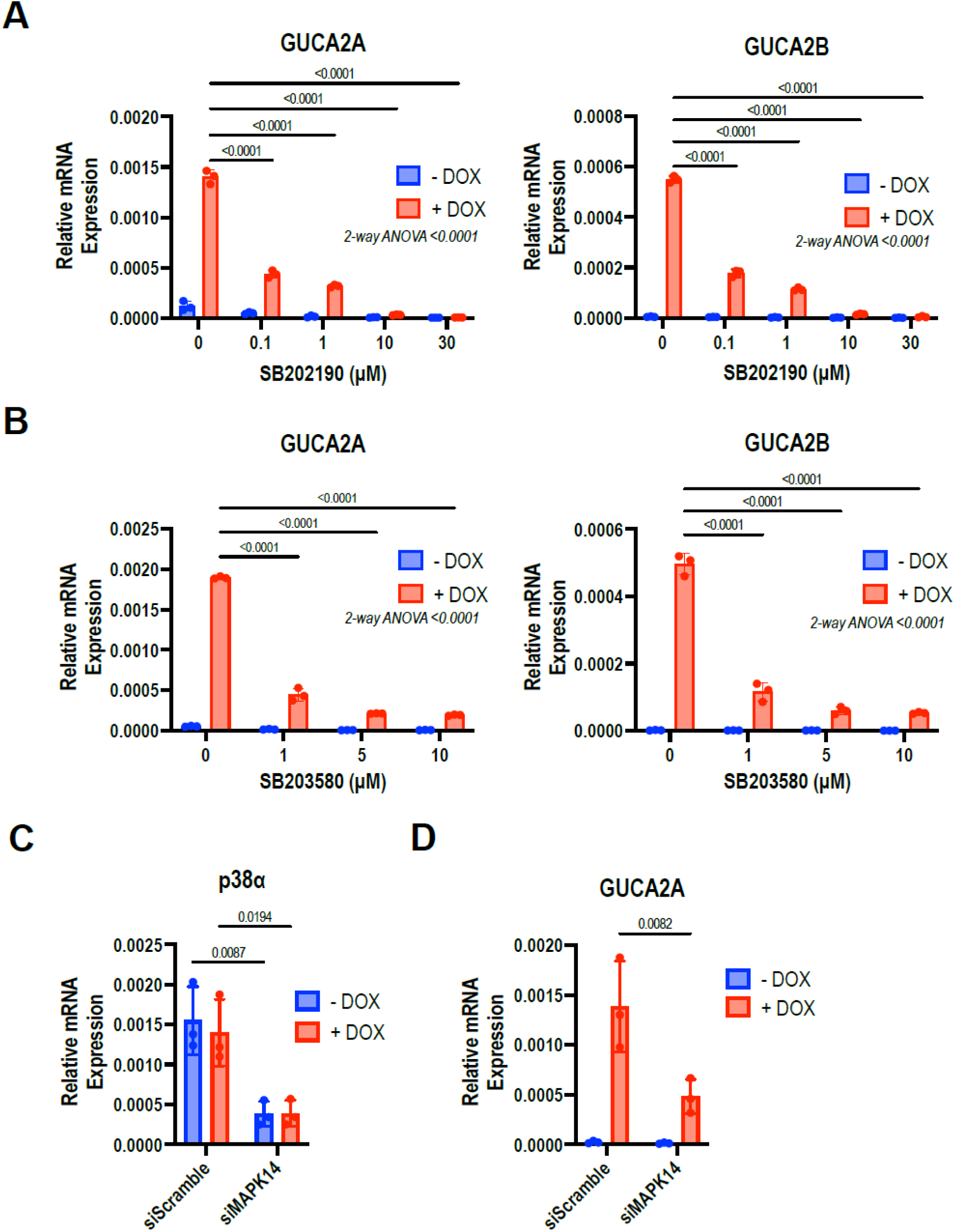
p38α/β inhibitors suppress GUCA2A and GUCA2B mRNA expression. (A) The p38α/β inhibitor SB202190 suppressed GUCA2A and GUCA2B mRNA expression in LS174T (DNTCF) cells. (B) The p38α/β inhibitor SB203580 suppressed GUCA2A and GUCA2B mRNA expression in a concentration-dependent fashion in LS174T (DNTCF) cells. (C) p38α expression was reduced by siRNA knockdown in LS174T (DNTCF) cells. (D) siRNA knockdown of p38α reduced GUCA2A mRNA expression in LS174T (DNTCF) cells following silencing of APC/β-catenin/TCF signaling. Statistics were calculated using a two-way ANOVA with Tukey’s multiple comparisons test. Error bars represent the standard deviation (SD).

**Supplemental Figure 3.**
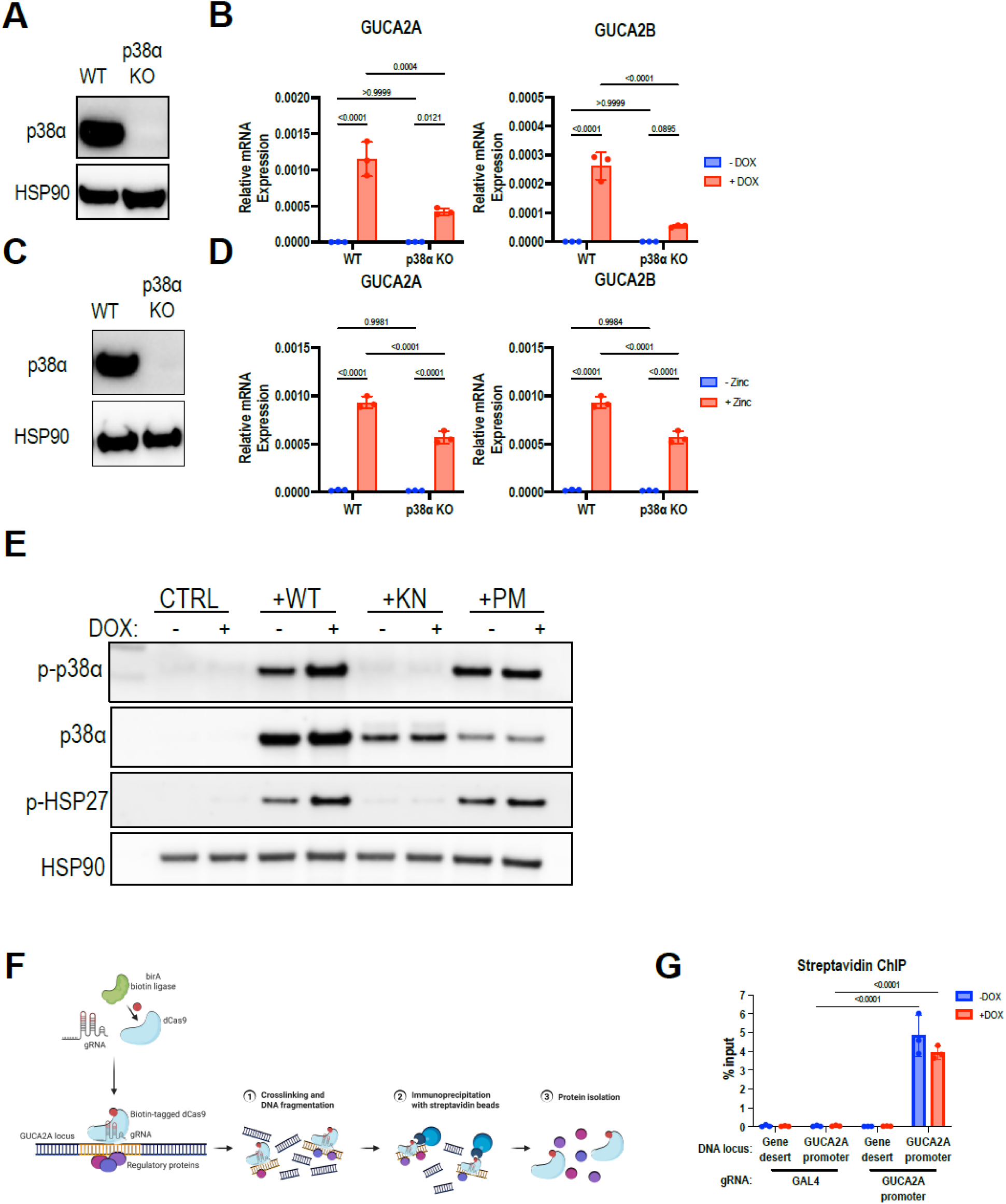
p38α KO reduces GUCA2A and GUCA2B mRNA expression in the absence of APC/β-catenin/TCF signaling. (A) p38α knockout in LS174T (siBC) results in (B) reduced GUCA2A and GUCA2B mRNA expression. (C) p38α knockout in HT29 (APC) cells results in (D) reduced GUCA2A and GUCA2B mRNA expression. (E) Exogenous p38α constructs (wildtype, WT; kinase-null, KN; phosphomimetic, PM) expressed in p38α knockout cells. (F) Schematic of the CRISPR affinity purification in situ of regulatory elements (CAPTURE) workflow. A catalytically dead Cas9 (dCas9) fused to a biotin acceptor peptide is directed to the target genomic locus by a guide RNA (gRNA). In the presence of the prokaryotic biotin ligase BirA, dCas9 is biotinylated, and protein complexes associated with the targeted locus are chemically crosslinked. Streptavidin affinity purification enriches locus-associated proteins for downstream proteomic analysis. (G) Using a *GUCA2A* promoter-targeting gRNA with birA-biotinylated dCas9, the *GUCA2A* promoter locus is enriched by streptavidin-targeted ChIP. There is no enrichment of a negative control gene desert in chromosome 12, and a non-targeting GAL4 gRNA does not enrich either locus. Statistics were calculated using a two-way ANOVA with Tukey’s multiple comparisons test. Error bars represent the standard deviation (SD).

**Supplemental Figure 4.**
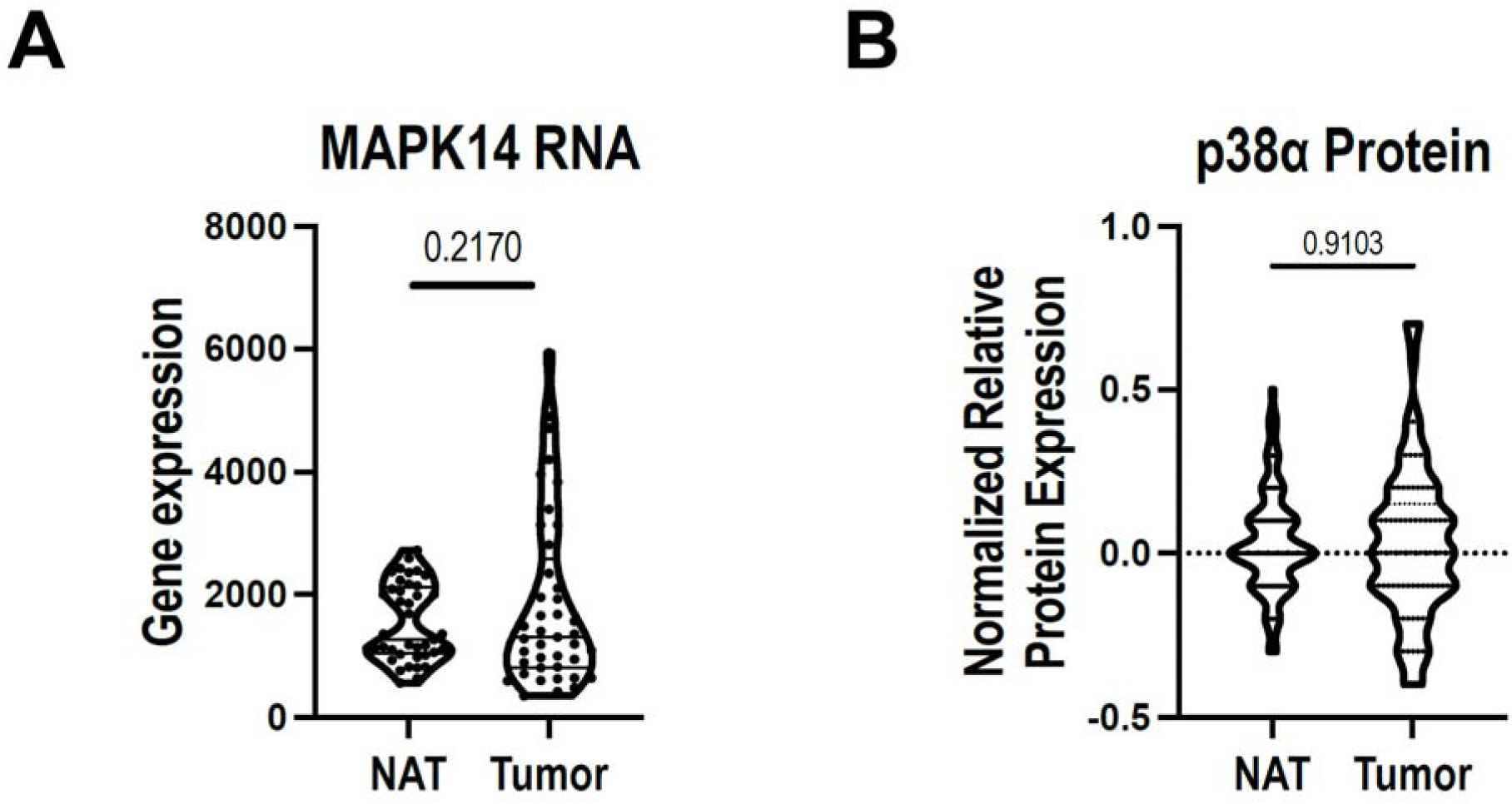
p38α expression is comparable in colorectal tumors and NAT. (A) mRNA of p38α in NAT and colorectal tumors from the TCGA COAD dataset. (B) p38α protein expression in NAT and colorectal tumors from the CPTAC colorectal cancer dataset. No significant differences in p38α mRNA or protein expression were observed between normal and tumor tissues. Statistics were calculated using an unpaired Welch’s *t* test.

**Supplemental Figure 5.**
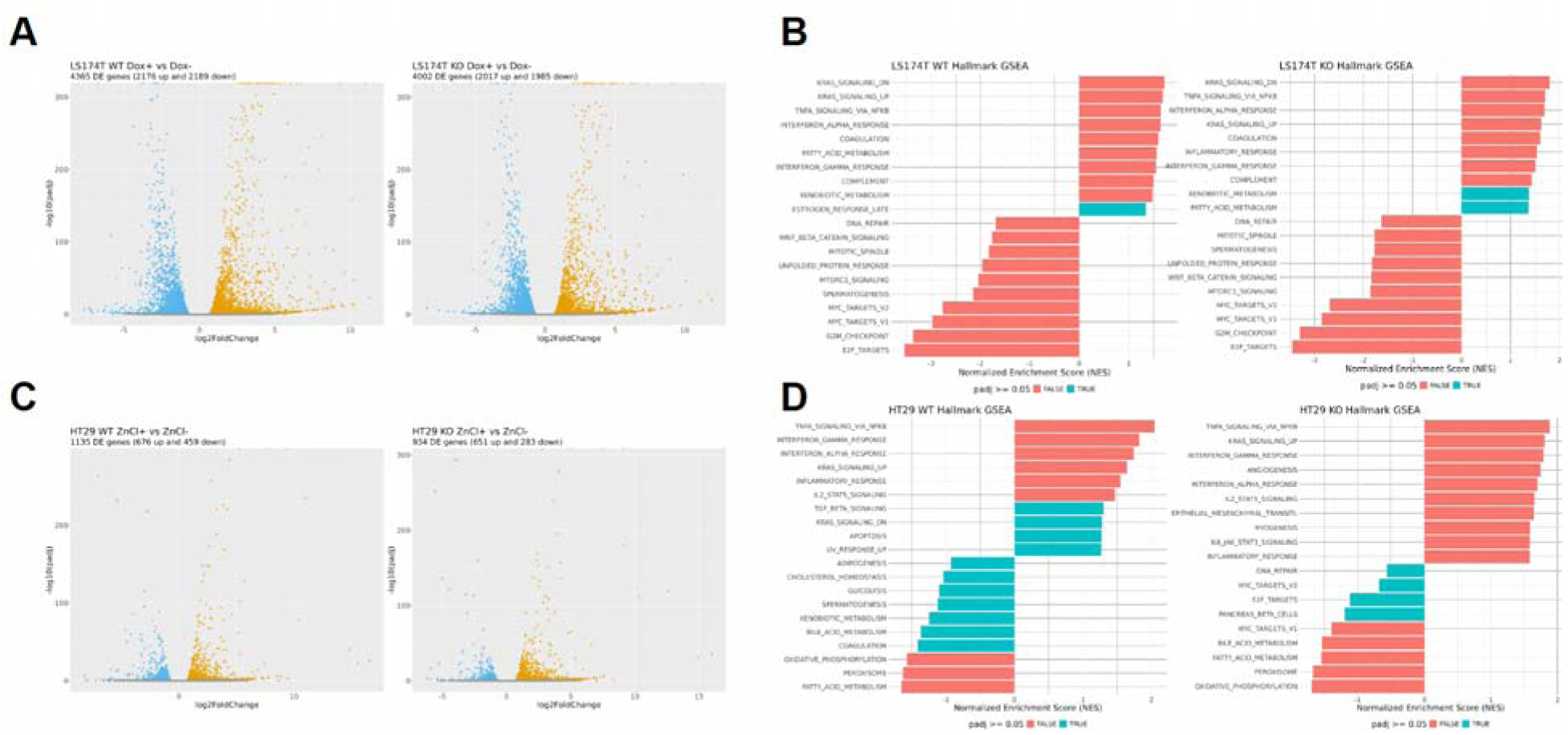
p38α-dependent signatures in LS174T (siBC) and HT29 (APC) cells. (A) Volcano plots of genes silenced or activated by APC/β-catenin/TCF signaling in LS174T (siBC) cells with or without p38α. (B) Significant hallmark pathways in LS174T (siBC) p38α WT and p38α KO cells. p_adj_<0.05 is denoted in red. (C) Volcano plots of genes silenced or activated by APC/β-catenin/TCF signaling in HT29 (APC) cells with or without p38α. (D) Significant hallmark pathways in HT29 (APC) p38α WT and p38α KO cells. p_adj_<0.05 is denoted in red.

## Supplemental Tables

Supplemental Table 1: Materials and reagents.

Supplemental Table 2: Cell line RNA-seq and pathway enrichment results.

Supplemental Table 3: AME analysis of putative promoters in the p38-dependent gene signature.

Supplemental Table 4: GSEA of the p38-dependent gene signature in the TCGA-COAD cohort.

## References

1. Zhang, L. & Shay, J. W. Multiple Roles of APC and its Therapeutic Implications in Colorectal Cancer. JNCI J. Natl. Cancer Inst. 109, djw332 (2017).

2. Rappaport, J. A. et al. A β-Catenin-TCF-Sensitive Locus Control Region Mediates GUCY2C Ligand Loss in Colorectal Cancer. Cell. Mol. Gastroenterol. Hepatol. 13, 1276–1296 (2022).

3. Harmston, N. et al. Widespread Repression of Gene Expression in Cancer by a Wnt/β-Catenin/MAPK Pathway. Cancer Res. 81, 464–475 (2021).

4. Osaki, L. H. & Gama, P. MAPKs and Signal Transduction in the Control of Gastrointestinal Epithelial Cell Proliferation and Differentiation. Int. J. Mol. Sci. 14, 10143–10161 (2013).

5. Cuadrado, A. & Nebreda, A., Cuadrado A, Nebreda ARMechanisms and functions of p38 MAPK signalling. Biochem J 429: 403–417. Biochem. J. 429, 403–17 (2010).

6. Phan, T., Zhang, X. H., Rosen, S. & Melstrom, L. G. P38 kinase in gastrointestinal cancers. Cancer Gene Ther. 30, 1181–1189 (2023).

7. Houde, M. et al. Intestinal Epithelial Cell Differentiation Involves Activation of p38 Mitogen-activated Protein Kinase That Regulates the Homeobox Transcription Factor CDX2 *. J. Biol. Chem. 276, 21885–21894 (2001).

8. Feng, Y. J. & Li, Y. Y. The role of p38 mitogen-activated protein kinase in the pathogenesis of inflammatory bowel disease. J. Dig. Dis. 12, 327–332 (2011).

9. Wakeman, D. et al. p38 MAPK regulates Bax activity and apoptosis in enterocytes at baseline and after intestinal resection | American Journal of Physiology-Gastrointestinal and Liver Physiology. https://journals.physiology.org/doi/full/10.1152/ajpgi.00485.2011.

10. Wakeman, D. et al. Deletion of p38-alpha mitogen-activated protein kinase within the intestinal epithelium promotes colon tumorigenesis. Surgery 152, 286–293 (2012).

11. Currie, M. G. et al. Guanylin: an endogenous activator of intestinal guanylate cyclase. Proc. Natl. Acad. Sci. U. S. A. 89, 947–951 (1992).

12. Hamra, F. K. et al. Uroguanylin: structure and activity of a second endogenous peptide that stimulates intestinal guanylate cyclase. Proc. Natl. Acad. Sci. U. S. A. 90, 10464–10468 (1993).

13. Field, M., Graf, L. H., Laird, W. J. & Smith, P. L. Heat-stable enterotoxin of Escherichia coli: in vitro effects on guanylate cyclase activity, cyclic GMP concentration, and ion transport in small intestine. Proc. Natl. Acad. Sci. U. S. A. 75, 2800–2804 (1978).

14. Hughes, J. M., Murad, F., Chang, B. & Guerrant, R. L. Role of cyclic GMP in the action of heat-stable enterotoxin of Escherichia coli. Nature 271, 755–756 (1978).

15. Lin, J. E. et al. GUCY2C Opposes Systemic Genotoxic Tumorigenesis by Regulating AKT-Dependent Intestinal Barrier Integrity. PLOS ONE 7, e31686 (2012).

16. Lin, J. E. et al. The Hormone Receptor GUCY2C Suppresses Intestinal Tumor Formation by Inhibiting AKT Signaling. Gastroenterology 138, 241–254 (2010).

17. Lin, J. E. et al. Obesity-induced colorectal cancer is driven by caloric silencing of the guanylin-GUCY2C paracrine signaling axis. Cancer Res. 76, 339–346 (2016).

18. Li, P. et al. Guanylyl cyclase C suppresses intestinal tumorigenesis by restricting proliferation and maintaining genomic integrity. Gastroenterology 133, 599–607 (2007).

19. Basu, N., Saha, S., Khan, I., Ramachandra, S. G. & Visweswariah, S. S. Intestinal Cell Proliferation and Senescence Are Regulated by Receptor Guanylyl Cyclase C and p21. J. Biol. Chem. 289, 581–593 (2014).

20. Shailubhai, K. et al. Uroguanylin treatment suppresses polyp formation in the Apc(Min/+) mouse and induces apoptosis in human colon adenocarcinoma cells via cyclic GMP. Cancer Res. 60, 5151–5157 (2000).

21. Wilson, C. et al. The paracrine hormone for the GUCY2C tumor suppressor, guanylin, is universally lost in colorectal cancer. Cancer Epidemiol. Biomark. Prev. Publ. Am. Assoc. Cancer Res. Cosponsored Am. Soc. Prev. Oncol. 23, 2328–2337 (2014).

22. Steinbrecher, K. A. et al. Expression of guanylin is downregulated in mouse and human intestinal adenomas. Biochem. Biophys. Res. Commun. 273, 225–230 (2000).

23. Pattison, A. M. et al. Silencing the intestinal GUCY2C tumor suppressor axis requires APC loss of heterozygosity. Cancer Biol. Ther. 21, 799–805 (2020).

24. Blomain, E. S. et al. APC-β-catenin-TCF signaling silences the intestinal guanylin-GUCY2C tumor suppressor axis. Cancer Biol. Ther. 21, 441–451 (2020).

25. van de Wetering, M. et al. The beta-catenin/TCF-4 complex imposes a crypt progenitor phenotype on colorectal cancer cells. Cell 111, 241–250 (2002).

26. Basken, J. et al. Specificity of Phosphorylation Responses to Mitogen Activated Protein (MAP) Kinase Pathway Inhibitors in Melanoma Cells. Mol. Cell. Proteomics MCP 17, 550–564 (2018).

27. Morin, P. J., Vogelstein, B. & Kinzler, K. W. Apoptosis and APC in colorectal tumorigenesis. Proc. Natl. Acad. Sci. U. S. A. 93, 7950–7954 (1996).

28. van de Wetering, M. et al. Specific inhibition of gene expression using a stably integrated, inducible small-interfering-RNA vector. EMBO Rep. 4, 609–615 (2003).

29. Han, J., Wu, J. & Silke, J. An overview of mammalian p38 mitogen-activated protein kinases, central regulators of cell stress and receptor signaling. F1000Research 9, F1000 Faculty Rev-653 (2020).

30. Freshney, N. W. et al. Interleukin-1 activates a novel protein kinase cascade that results in the phosphorylation of Hsp27. Cell 78, 1039–1049 (1994).

31. Michels, B. E. et al. Human colon organoids reveal distinct physiologic and oncogenic Wnt responses. J. Exp. Med. 216, 704–720 (2019).

32. Brenna, Ø. et al. Cellular localization of guanylin and uroguanylin mRNAs in human and rat duodenal and colonic mucosa. Cell Tissue Res. 365, 331–341 (2016).

33. Ikpa, P. T. et al. Guanylin and uroguanylin are produced by mouse intestinal epithelial cells of columnar and secretory lineage. Histochem. Cell Biol. 146, 445–455 (2016).

34. Qian, X., Prabhakar, S., Nandi, A., Visweswariah, S. S. & Goy, M. F. Expression of GC-C, a receptor-guanylate cyclase, and its endogenous ligands uroguanylin and guanylin along the rostrocaudal axis of the intestine. Endocrinology 141, 3210–3224 (2000).

35. Enslen, H., Raingeaud, J. & Davis, R. J. Selective activation of p38 mitogen-activated protein (MAP) kinase isoforms by the MAP kinase kinases MKK3 and MKK6. J. Biol. Chem. 273, 1741–1748 (1998).

36. Diskin, R., Askari, N., Capone, R., Engelberg, D. & Livnah, O. Active mutants of the human p38alpha mitogen-activated protein kinase. J. Biol. Chem. 279, 47040– 47049 (2004).

37. Liu, X., et al. *In Situ* Capture of Chromatin Interactions by Biotinylated dCas9. Cell 170, 1028–1043.e19 (2017).

38. Ferreiro, I. et al. Whole genome analysis of p38 SAPK-mediated gene expression upon stress. BMC Genomics 11, 144 (2010).

39. Cuenda, A. & Rousseau, S. p38 MAP-Kinases pathway regulation, function and role in human diseases. Biochim. Biophys. Acta BBA - Mol. Cell Res. 1773, 1358–1375 (2007).

40. Vasaikar, S. et al. Proteogenomic analysis of human colon cancer reveals new therapeutic opportunities. Cell 177, 1035–1049.e19 (2019).

41. Raingeaud, J., Whitmarsh, A. J., Barrett, T., Dérijard, B. & Davis, R. J. MKK3- and MKK6-regulated gene expression is mediated by the p38 mitogen-activated protein kinase signal transduction pathway. Mol. Cell. Biol. 16, 1247–1255 (1996).

42. Wang, P.-N. et al. Downregulation of phosphorylated MKK4 is associated with a poor prognosis in colorectal cancer patients. Oncotarget 8, 34352–34361 (2017).

43. Bailey, T. L., Johnson, J., Grant, C. E. & Noble, W. S. The MEME Suite. Nucleic Acids Res. 43, W39–W49 (2015).

44. Zarubin, T. & Han, J. Activation and signaling of the p38 MAP kinase pathway. Cell Res. 15, 11–18 (2005).

45. Han, J., Wu, J. & Silke, J. An overview of mammalian p38 mitogen-activated protein kinases, central regulators of cell stress and receptor signaling. F1000Research 9, F1000 Faculty Rev-653 (2020).

46. Corre, I., Paris, F. & Huot, J. The p38 pathway a major pleiotropic cascade that transduces stress and metastatic signals in endothelial cells. Oncotarget 8, 55684– 55714 (2017).

47. Nyati, K. K. et al. TLR4-induced NF-κB and MAPK signaling regulate the IL-6 mRNA stabilizing protein Arid5a. Nucleic Acids Res. 45, 2687–2703 (2017).

48. Xu, K. & Shu, H.-K. G. EGFR activation results in enhanced cyclooxygenase-2 expression through p38 mitogen-activated protein kinase-dependent activation of the Sp1/Sp3 transcription factors in human gliomas. Cancer Res. 67, 6121–6129 (2007).

49. Hermeling, J. C. et al. Mitochondria-originated redox signalling regulates KLF-1 to promote longevity in Caenorhabditis elegans. Redox Biol. 58, 102533 (2022).

50. del Barco Barrantes, I., Coya, J. M., Maina, F., Arthur, J. S. C. & Nebreda, A. R. Genetic analysis of specific and redundant roles for p38alpha and p38beta MAPKs during mouse development. Proc. Natl. Acad. Sci. U. S. A. 108, 12764–12769 (2011).

51. Warr, N. et al. Gadd45γ and Map3k4 Interactions Regulate Mouse Testis Determination via p38 MAPK-Mediated Control of Sry Expression. Dev. Cell 23, 1020– 1031 (2012).

52. Furlan, A. et al. Met acts through Abl to regulate p53 transcriptional outcomes and cell survival in the developing liver. J. Hepatol. 57, 1292–1298 (2012).

53. Saba-El-Leil, M. K., Frémin, C. & Meloche, S. Redundancy in the World of MAP Kinases: All for One. Front. Cell Dev. Biol. 4, (2016).

54. Llopis, A. et al. The stress-activated protein kinases p38α/β and JNK1/2 cooperate with Chk1 to inhibit mitotic entry upon DNA replication arrest. Cell Cycle 11, 3627–3637 (2012).

55. Hayakawa, M. et al. Loss of Functionally Redundant p38 Isoforms in T Cells Enhances Regulatory T Cell Induction *. J. Biol. Chem. 292, 1762–1772 (2017).

56. Qi, X. et al. p38alpha antagonizes p38gamma activity through c-Jun-dependent ubiquitin-proteasome pathways in regulating Ras transformation and stress response. J. Biol. Chem. 282, 31398–31408 (2007).

57. Yin, N. et al. p38γ MAPK is required for inflammation-associated colon tumorigenesis. Oncogene 35, 1039–1048 (2016).

58. Bikkavilli, R. K., Feigin, M. E. & Malbon, C. C. p38 mitogen-activated protein kinase regulates canonical Wnt–β-catenin signaling by inactivation of GSK3β. J. Cell Sci. 121, 3598–3607 (2008).

59. Chang, J. et al. Noncanonical Wnt-4 signaling enhances bone regeneration of mesenchymal stem cells in craniofacial defects through activation of p38 MAPK. J. Biol. Chem. 282, 30938–30948 (2007).

60. Thornton, T. M. et al. Phosphorylation by p38 MAPK as an alternative pathway for GSK3beta inactivation. Science 320, 667–670 (2008).

61. Lepore Signorile, M., et al. Uncoupling p38α nuclear and cytoplasmic functions and identification of two p38α phosphorylation sites on β-catenin: implications for the Wnt signaling pathway in CRC models. Cell Biosci. 13, 223 (2023).

62. Lepore Signorile, M., et al. Pharmacological targeting of the novel β-catenin chromatin-associated kinase p38α in colorectal cancer stem cell tumorspheres and organoids. Cell Death Dis. 12, 1–18 (2021).

63. Lepore Signorile, M., et al. Uncoupling p38α nuclear and cytoplasmic functions and identification of two p38α phosphorylation sites on β-catenin: implications for the Wnt signaling pathway in CRC models. Cell Biosci. 13, 223 (2023).

64. Basu, N., Saha, S., Khan, I., Ramachandra, S. G. & Visweswariah, S. S. Intestinal Cell Proliferation and Senescence Are Regulated by Receptor Guanylyl Cyclase C and p21. J. Biol. Chem. 289, 581–593 (2014).

65. Gupta, J. et al. Dual function of p38α MAPK in colon cancer: suppression of colitis-associated tumor initiation but requirement for cancer cell survival. Cancer Cell 25, 484–500 (2014).

66. Brancho, D. et al. Mechanism of p38 MAP kinase activation in vivo. Genes Dev. 17, 1969–1978 (2003).

67. del Reino, P. et al. Pro-Oncogenic Role of Alternative p38 Mitogen-Activated Protein Kinases p38γ and p38δ, Linking Inflammation and Cancer in Colitis-Associated Colon Cancer. Cancer Res. 74, 6150–6160 (2014).

68. Chiacchiera, F. et al. p38alpha blockade inhibits colorectal cancer growth in vivo by inducing a switch from HIF1alpha-to FoxO-dependent transcription. Cell Death Differ. 16, 1203–1214 (2009).

69. Chiacchiera, F. et al. Blocking p38/ERK crosstalk affects colorectal cancer growth by inducing apoptosis in vitro and in preclinical mouse models. Cancer Lett. 324, 98–108 (2012).

70. Mölder, F. et al. Sustainable data analysis with Snakemake. F1000Research 10, 33 (2021).

71. Bolger, A. M., Lohse, M. & Usadel, B. Trimmomatic: a flexible trimmer for Illumina sequence data. Bioinformatics 30, 2114–2120 (2014).

72. Dobin, A. et al. STAR: ultrafast universal RNA-seq aligner. Bioinformatics 29, 15– 21 (2013).

73. Li, B. & Dewey, C. N. RSEM: accurate transcript quantification from RNA-Seq data with or without a reference genome. BMC Bioinformatics 12, 323 (2011).

74. Love, M. I., Huber, W. & Anders, S. Moderated estimation of fold change and dispersion for RNA-seq data with DESeq2. Genome Biol. 15, 550 (2014).

75. Subramanian, A. et al. Gene set enrichment analysis: a knowledge-based approach for interpreting genome-wide expression profiles. Proc. Natl. Acad. Sci. U. S. A. 102, 15545–15550 (2005).

76. Liberzon, A. et al. The Molecular Signatures Database (MSigDB) hallmark gene set collection. Cell Syst. 1, 417–425 (2015).

77. Korotkevich, G. et al. Fast gene set enrichment analysis. 060012 Preprint at 10.1101/060012 (2021).

78. Bugter, J. M., Fenderico, N. & Maurice, M. M. Mutations and mechanisms of WNT pathway tumour suppressors in cancer. Nat. Rev. Cancer 21, 5–21 (2021).

79. Baybutt, T. R. et al. CD8α Structural Domains Enhance GUCY2C CAR-T Cell Efficacy. Cancer Biol. Ther. 25, 2398801.

80. Casper, J. et al. The UCSC Genome Browser database: 2026 update. Nucleic Acids Res. 54, D1331–D1335 (2026).

81. Khan, A. et al. JASPAR 2018: update of the open-access database of transcription factor binding profiles and its web framework. Nucleic Acids Res. 46, D1284 (2018).

